# Modulation of Recovery from Neonatal Hyperoxic Lung Injury by Sex as a Biological Variable

**DOI:** 10.1101/2023.08.09.552532

**Authors:** Abiud Cantu, Manuel Cantu Gutierrez, Xiaoyu Dong, Connor Leek, Montserrat Anguera, Krithika Lingappan

**Author notes:** Corresponding Author: Krithika Lingappan, MD, MS. Ph.D. Address: 3401 Civic center Blvd, Philadelphia, PA, 19104.

## Abstract

Recovery from lung injury during the neonatal period requires the orchestration of many biological pathways. The modulation of such pathways can drive the developing lung towards proper repair or persistent maldevelopment that can lead to a disease phenotype. Sex as a biological variable can regulate these pathways differently in the male and female lung exposed to neonatal hyperoxia. In this study, we assessed the contribution of cellular diversity in the male and female neonatal lung following injury. Our objective was to investigate sex and cell-type specific transcriptional changes that drive repair or persistent injury in the neonatal lung and delineate the alterations in the immune-endothelial cell communication networks using single cell RNA sequencing (sc-RNAseq) in a murine model of hyperoxic injury. We generated transcriptional profiles of >55,000 cells isolated from the lungs of postnatal day 1 (PND 1) and postnatal day 21 (PND 21) neonatal male and female C57BL/6 mice exposed to 95% FiO_2_ between PND 1-5 (saccular stage of lung development). We show the presence of sex-based differences in the transcriptional states of lung endothelial and immune cells at PND 1 and PND 21. Furthermore, we demonstrate that biological sex significantly influences the response to injury, with a greater number of differentially expressed genes showing sex-specific patterns than those shared between male and female lungs. Pseudotime trajectory analysis highlighted genes needed for lung development that were altered by hyperoxia. Finally, we show intercellular communication between endothelial and immune cells at saccular and alveolar stages of lung development with sex-based biases in the crosstalk and identify novel ligand-receptor pairs. Our findings provide valuable insights into the cell diversity, transcriptional state, developmental trajectory, and cell-cell communication underlying neonatal lung injury, with implications for understanding lung development and possible therapeutic interventions while highlighting the crucial role of sex as a biological variable.

## Introduction

The male disadvantage for neonatal mortality and major morbidities in preterm neonates is well known [1–4]. Sex-chromosome-based, hormonal, or epigenetic mechanisms may modulate the male susceptibility or the female resilience [5]. Susceptibility to diseases and the subsequent repair and recovery may differ between the sexes due to the sex-based differences at baseline or pathways activated or inhibited in response to the injurious stimuli.

Poor lung health in premature neonates is a major factor underlying an inferior quality of life, high economic costs related to health care and a greater risk of developing adult-onset chronic lung diseases [6,7]. Respiratory morbidity including the development of bronchopulmonary dysplasia (BPD) is common in preterm neonates with long term impact. Even in the post-surfactant era, extremely premature male neonates (born between 24 and 26 weeks of gestation) display a significantly increased risk of developing BPD [8]. The incidence of respiratory distress syndrome (RDS), BPD, and moderate to severe BPD was significantly increased in preterm males after adjusting for multiple confounding factors [9]. Similarly, the need for respiratory support, respiratory medications and home oxygen use was higher in males [10,11]. Male sex was an independent risk factor for tracheostomy among preterm neonates and in infants who received a tracheostomy, an independent predictor of mortality [12]. Hospital readmissions and outpatient visits due to respiratory issues are also increased in male premature neonates [13]. Male sex is thus an independent predictor for the development of BPD and the subsequent morbidities related to lung disease in premature infants, but the underlying mechanisms behind these sexually dimorphic outcomes are unknown.

Animal models for human BPD use several approaches to replicate the major findings of alveolar simplification and abnormal vascular remodeling in the murine lung including postnatal hyperoxia exposure in the term mouse [14,15]. Exposure to hyperoxia and the resulting oxidative stress to the developing lung contributes to the pathophysiology of this disease. The murine lung is in the saccular stage of lung development from birth to postnatal day (PND4-5), which is equivalent to 26-36 weeks in human preterm neonates [14,16]. Most preterm neonates are exposed to varying degrees of hyperoxia in the neonatal intensive care unit during this period, with the sicker neonates receiving the highest concentrations of oxygen support.

Our lab and others have reported sex-specific differences in alveolar and vascular development in neonatal mice using the postnatal hyperoxia model [17,18]. Furthermore, we have highlighted differences involving epigenetic mechanisms, micro-RNA mediated effects and the role of sex-chromosome and gonadal hormones in this model all of which show striking sex-specific differences [19–22]. However, the sex-specific differences in individual lung cell sub populations at single-cell resolution immediately after birth at PND1 and after hyperoxic injury at PND21 (alveolar stage of lung development) are not known. In this investigation, using sc-RNA Seq, we tested the hypothesis that sex as a biological variable will modulate biological pathways differently in the male and female lung exposed to neonatal hyperoxia.

## MATERIALS AND METHODS

### Mice

All animal experiments were performed under an approved protocol by the IACUC at the Children’s Hospital of Philadelphia. Timed pregnant C57BL/6N WT mice were obtained from Charles River Laboratories (Wilmington). The sex in neonatal mouse pups was determined by both the anogenital distance and pigmentation in the anogenital region method. In neonatal male mice, a pigmented spot on the scrotum is visible to the naked eye from postnatal day (PND) 1, whereas female pups lack visible pigmentation in the anogenital region. This was also verified by the expression of Y-chromosome transcript abundance in male samples.

### Mouse model of BPD

An arrest of alveolarization was induced in mouse pups by exposure to hyperoxia (95% O2), as described previously [23]. Mouse pups from multiple litters were pooled before being randomly and equally redistributed to two groups, one group exposed to room air (21% O2) and the other group exposed to hyperoxia (95% O2), within 12 h of birth for 5 days. The dams were rotated between air- and hyperoxia-exposed litters every 24h.

### Lung Isolation for Single Cell Sequencing

Mice were euthanized on postnatal day (PND) 1 and 21 with i.p. pentobarbital. The right ventricle was perfused with ice cold PBS and the lungs were harvested quickly and homogenized and dissociated using the Miltenyi MACS lung dissociation kit (Miltenyi Biotec, cat # 130-095-927), using the GentleMacs program m-Lung-1 on the AUTO MaCs. The lungs were incubated for 20min at 37 ◦C under continuous rotation and then run on the gentleMACS m-Lung-2 program on the gentleMACS (Miltenyi Biotec, Cat #130-095-937). This was followed by brief centrifugation and the passage of cells through the 70-uM filter, washed, centrifuged again and the pellet was treated with ACK lysis buffer, centrifuged, and resuspended in sorting buffer and passed through an 40uM filter. Live cells were sorted and subjected to single-cell RNA sequencing as described below.

### scRNA-Seq Library Preparation and Sequencing

Single cell Gene Expression Library was prepared according to Chromium Single Cell Gene Expression 3v3.1 kit (10x Genomics). In brief, single cells, reverse transcription (RT) reagents, Gel Beads containing barcoded oligonucleotides, and oil were loaded on a Chromium controller (10x Genomics) to generate single cell GEMS (Gel Beads-In-Emulsions) where full length cDNA was synthesized and barcoded for each single cell. Subsequently the GEMS are broken and cDNA from each single cell are pooled. Following cleanup using Dynabeads MyOne Silane Beads, cDNA is amplified by PCR. The amplified product is fragmented to optimal size before end-repair, A-tailing, and adaptor ligation. Final library was generated by amplification. 10X Genomics Cell Ranger [version] ‘cellranger count’ was used to perform alignment, filtering, barcode counting, and UMI counting. The raw data has been uploaded to NCBI GEO; GSE237944.

### Sequencing, data processing, quality control, integration, and cluster annotation

A Seurat [24] object was generated taking the 10x Genomics Cell Ranger 6.0.0 [25] output matrices using Read10X method for each of the sequenced samples. SCTransformV2 [26] and a GetClusters function (RunPCA, RunUMAP, RunTSNE, FindNeighbors, FindClusters) was run. SoupX version 1.6.1 [27] was run with a contamination factor of 5% to reduce mRNA background contamination. To get percent of mitochondrial genes in each cell we used PercentageFeatureSet. Resulting cells were filtered using >= 500 molecular identifiers (nUMI >= 500), removing upper number of UMI outliers (nUMI <= outlier_threshold_nUMI), >250 detectable genes (nGene >= 250), removing upper number of gene outliers (nGene<= outlier_threshold_nGene), log10GenesPerUMI > 0.80, < 5% of transcripts coming from mitochondrial genes (mitoRatio < 0.05). Only genes expressed in more than 10 cells were kept. The genes ‘*Gm42418*’, ‘*S100a8*’, ‘*S100a9*’ were removed from Seurat object. We ran SCTransformV2 and GetClusters (RunPCA, RunUMAP, RunTSNE, FindNeighbors, FindClusters) again with a resolution of 0.6. Doublets were removed using DoubletFinder version 2.0.3 [28] using the paramSweep_v3, summarizeSweep, find.pK, doubletFinder_v3 functions. GetClusters was run again post filtering of doublets. We aggregated the individual samples using SelectIntegrationFeatures with number of features to return set to 3000, PrepSCTIntegration, FindIntegrationAnchors with sample 1 as reference (female room air for PND 1 samples and female hyperoxia for PND 21 samples). IntegrateData using SCT as normalization method was then applied. GetClusters was used again with a resolution of 0.6. For cell type assignment, PrepSCTFindMarkers was ran with default parameters and FindMarkers set to Only return positive markers using wilcoxon test and using the SCT assay. Differential expression between conditions were found using Seurat’s FindMarkers function. Genes detected in 25% of cells with an adjusted p-value of < 0.05 and a log2fold change of > 0.3 were reported as significant in each comparison. Significant genes were then processed using enrichR[29] (version 3.0) along fgsea [30] to find enriched biological pathways present. Pseudotime analysis was done using Monocle3 [31]. Cell-cell communication was analyzed using Cellchat [32] on room air and hyperoxia samples separately with the software’s default parameters. The complete code used in this manuscript can be found in https://github.com/Abiud/Lingappan2023_Modulation_of_Recovery_pnd1_21_scRNASe <u>q</u>.

### *In Situ* Hybridization

In situ hybridization was performed using the RNAscope Multiplex Fluorescent Reagent Kit v2 (323110, Advanced Cell Diagnostics, Hayward, CA, USA) with RNAscope 4-Plex Ancillary Kit for Multiplex Fluorescent V2 (232120, Advanced Cell Diagnostics, Hayward, CA, USA). Samples were treated with *Xist* (418281), *Tsc22d3* (448341), *Adgre1* (460651-C2), *Siglec F* (518461-C3), and *Pecam* (316721-C4) probes (Advanced Cell Diagnostics, Hayward, CA, USA) for 2 hours at 4°C. For each round, samples were run alongside a positive control slide treated with housekeeping genes *Polr2A, Ppib, Ubc, and Hprt* (321811) and a negative control (321831). Nuclei were counterstained with DAPI and autofluorescence was reduced using TrueBlack Plus Lipofuscin Autofluoresne Quencer 40X in DMSO (Biotum: 23014) for 30 minutes then washed in PBS. Slides were mounted in EverBrite TrueBlack® Hardset Mounting Medium (Biotium: 23017). Slides were imaged using Leica Bmi8 Thunder Imager microscope.

## RESULTS

### Developmental stage and sex modulate the cellular transcriptional state in the developing lung

To gain a comprehensive understanding of the sex specific transcriptional changes in the neonatal lung at PND 1 (saccular stage of lung development), PND 7 (early alveolar stage of lung development) and PND 21 (late alveolar stage of lung development), we collected lungs from male and female neonatal mice under room air conditions and processed them for single cell RNA sequencing (**Figure 1A**). The data from PND 7 was used from our previously published work [23]. More than 55,000 cell transcriptomes were profiled with cell clustering showing capture overlap between cells in both male and female lungs (**Figure 1B**). *Epcam+* epithelial, *Cldn5*+ endothelial, *Ptprc*+ immune and *Col1a2+* mesenchymal cells were identified (**Figure 1C**). We visualized the data with Uniform Manifold Approximation and Projection (UMAP) at PND 1 and PND 21 room air lungs (**Figure 1D-E**). Annotation of these cell sub-populations was done based on prior publications focused on the developing and adult murine lungs [23,33–37]. Cellular annotation markers and their distribution among different cellular sub-clusters were also labeled (**Supplemental Figure 1, Supplemental Table 1 and 2**). Certain cellular populations like transient macrophages and proliferating ECs were only seen at PND 1 and not at PND 21, pointing to a developmental stage specific function of these cells. This has been previously highlighted by other publications [33,34]. The number of cells sequenced at each time-point and their relative distribution by sex is shown in **Figure 1F**. Notably, our lung digestion protocols yielded more immune and endothelial cell populations. At PND 21, the yield of epithelial and mesenchymal cells was lower compared to the earlier time-points. For each of the lung cell sub-populations, we compared the transcriptional profile between male and female (maintained in room air) lung cells (**Figure 1G**). Markedly, gene-expression varies in these cells by sex at all three time-points. For example, alveolar macrophages show greatest number of differentially expressed genes between the male and female lung at PND 1 which then decrease with time. B cells on the other hand show the greatest differences at PND 7, while T cells at PND 21.

**Figure 1:**
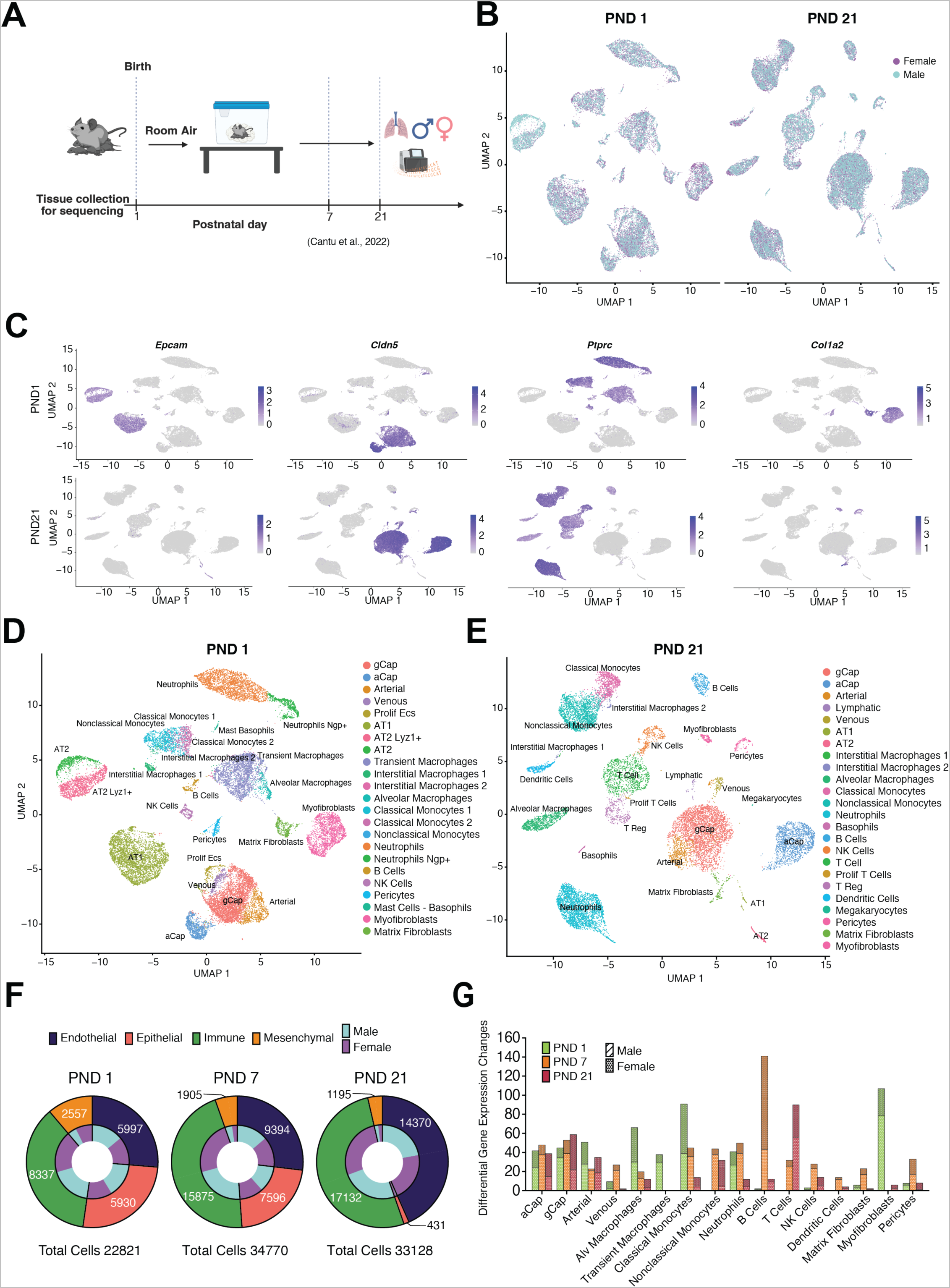
Developmental stage and sex modulates cellular transcriptional state in the developing lung. **(A)** Experimental design and timeline. Mouse pups (C57BL6) from multiple litters were pooled before being randomly and equally redistributed to two groups, one group exposed to room air (21% O2) and the other group exposed to hyperoxia (95% O2), within 12 h of birth for 5 days, and euthanized on PND 21. Lung from male and female lungs were also obtained at PND 1. Equal number of pooled viable lung cells from three mice were used for each group. **(B)** UMAP of sequenced cells labeled by sex (male or female). **(C)** UMAP overview of cell clusters identified based on gene expression. Epithelial (*Epcam+*), endothelial (*Cldn5+*), immune (*Ptprc+*) and mesenchymal (*Col1a2+*). **(D and E)** Annotation of lung cell sub-populations at PND 1 and PND 21. **(F)** The number of cells sequenced at each time-point and their relative distribution by sex. **(G)** Number of differentially expressed genes between male and female lung cells at three lung developmental stages PND 1 (saccular), PND 7 (early alveolar), and PND 21 (late alveolar).

### The lung microvascular endothelial cells show sex specific gene expression signatures during development

Sub clustering of lung endothelial cells identified distinct cell populations at PND 1 (**Figure 2A**) and PND 21 (**Figure 2B**). Lung capillary endothelial cells comprising of general capillaries (gCaps; Cap1) and aerocytes or alveolar capillaries (aCaps; Cap2) make up over 75% of the cells captured. A population of proliferating endothelial cells was identified at PND 1 but not at PND 21. A similar cluster was identified in our previous manuscript at PND 7 [23]. The cells in this cluster are positive for proliferation markers such as *Mki67* and *Top2a* (**Figure 2C**). Interestingly, these clusters are enriched for gCap markers such as *Kit*, *Itga6,* and *Npr3* and have low expression of aCap markers *Car4* and *Apln* (**Figure 2D**). Furthermore, the similarity between gCaps and proliferating ECs is highlighted when observing the correlation analysis of gene expression between clusters (**Figure 2E**). The highest correlation was within each cell type across time-points. Proliferating ECs showed a shared gene expression signature at PND 1 and PND 7 (**Figure 2F**, top) while also include time-point specific genes (**Figure 2F**, bottom heatmaps). Biological pathways enriched at PND 1 included interferon related pathways, while at PND 7: cell cycle, TGF-beta signaling, TNF signaling via NF-kappaB, endothelial cell migration, androgen and estrogen response pathways were enriched (**Figure 2G**).

**Figure 2:**
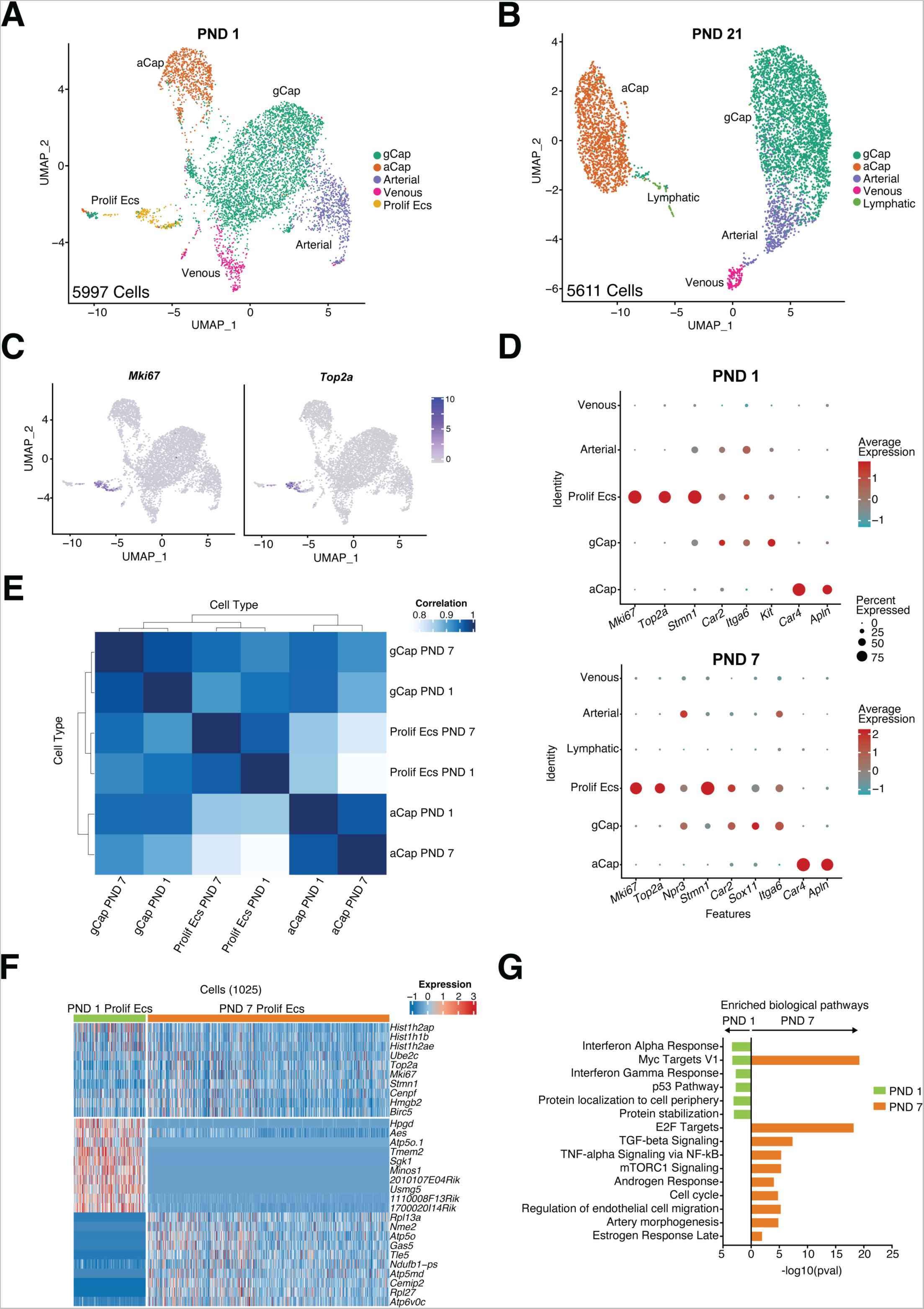
Proliferating endothelial cells are present early during postnatal lung development and show temporal changes in transcriptional state with increasing lung maturation. **(A-B)** UMAP of sequenced lung endothelial cells identified distinct clusters at PND 1 and PND 21. **(C)** Feature plots showing high expression of *Mki67* and *Top2a* in the proliferating endothelial cell subcluster. **(D)** Dot plot showing overlap of markers between gCaP and proliferating endothelial cells at PND 1 and PND 7. **(E)** Correlation of gene expression between proliferating ECs, gCaPs and aCaps. **(F)** Heatmap showing enriched genes in the proliferating ECs cluster at PND 1 and PND 7. **(G)** Enriched biological pathways common to and distinct in proliferating endothelial cells at PND 1 and PND 7. Number of cells sequenced are shown within parentheses.

Next, we wanted to identify the differences between male and female lung capillary endothelial cells at the PND 1 (saccular) and PND 21 (alveolar) stage of lung development. **Figures 3A-D** shows the differential expression in male vs. female, enriched pathways, and violin plots of selected genes in aCaps (**Supplemental Table 3**), while **Figures 3E-H** highlights the same in gCaps (**Supplemental Table 4**). As expected, X-chromosome localized genes such as *Xist* and *Ddx3x* are highly expressed in the female lung and Y-chromosome related genes such as *Ddx3y* are expressed in the male lung. We identified the biological pathways that were differentially regulated in the male and female aCaps and gCaps. Interestingly, in the female lung, both aCaps and gCaps at PND 1 enriched for pathways related to the immune system, cellular response related to cytokine stimulus and TNF-alpha signaling via NF-kappaB (**Figure 3B, F**). In contrast, in male lungs, immune mediated signaling particularly related to interferon response was enriched at PND 21 in both aCaps and gCaps (**Figure 3D, H**). These point to a close interaction between the developing endothelium and the lung immune cells that are active through different cellular signaling pathways in the male and female lung capillary endothelial cells at different stages of lung development.

**Figure 3:**
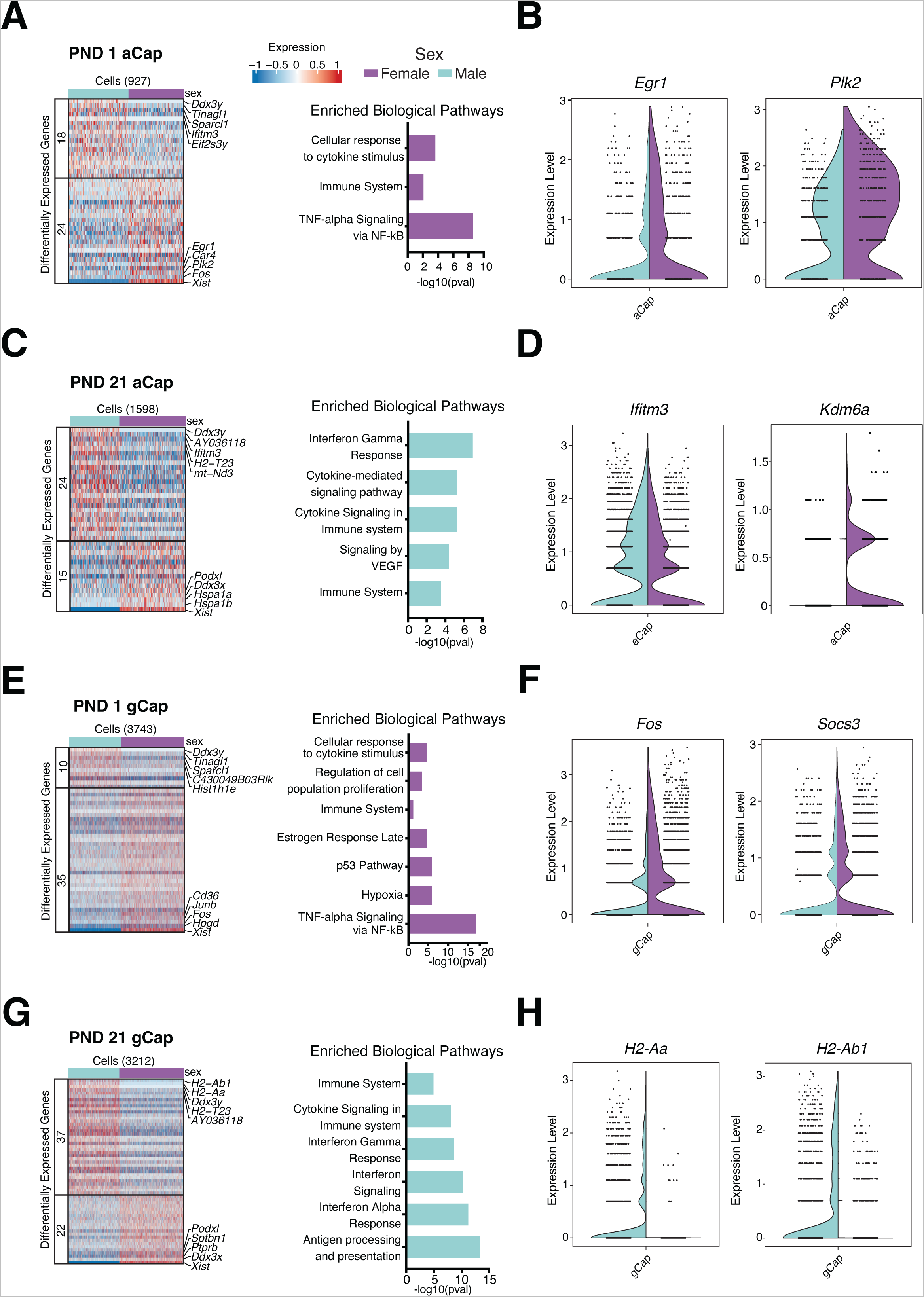
Sex-specific programming of biological pathways in the lung microvascular endothelial cells during lung development. **(A)** Heatmap of differentially expressed genes between male and female aCaps at PND 1 and **(C)** PND 21 along with enriched biological pathways. **(B)** Violin plots of differentially expressed genes in aCaps at PND 1 *Egr1* and *Plk2* and **(D)** PND 21 *Ifitm3* and *Kdm6a*. **(E)** Heatmap of differentially expressed genes between male and female gCaps at PND 1 and **(G)** PND 21 along with enriched biological pathways. **(F)** Violin plots of differentially expressed genes in gCaps at PND 1 *Fos* and *Socs3* and **(H)** PND 21 *H2-Aa* and *H2-Ab1*. Number of cells sequenced are shown within parentheses.

The transcription factor, early growth response-1 (*Egr1*) was increased in the PND 1 female aCaps (**Figure 3B**). *Egr1* increases angiogenesis and neovascularization [38], is induced by TNF-α [39,40], and EGR1 signaling axis plays a role in TNF-α-induced endothelial differentiation of human mesenchymal stem cells by increasing VEGFR2 expression [41]. The second highlighted gene, polo-like kinase 2 (*Plk2,* **Figure 3B**) plays a role in endothelial cell sprouting and angiogenesis with a loss of function resulting in decreased sprouting and migration in HUVECs [42] and loss of *Plk2* increases pulmonary fibrosis [43]. Interferon induced transmembrane protein 3 (*Ifitm3*) was increased in the male gCaps at PND 1 (**Figure 3D**) [44] and plays a role in decreasing virus-mediated pathology in the lung [45,46]. At PND 21, female aCaPs expressed *Kdm6a* (**Figure 3D**). *Kdm6a* encodes a histone demethylase that can modulate gene expression through epigenetic changes and modulates cell fate [47]. *Kdm6a* is located on the X-chromosome, is considered an X-chromosome inactivation (XCI) escapee gene, and is reliably expressed at higher levels in XX than XY cells in mice and humans [48–50]. Female gCaPs at PND 1 showed higher expression of Fos (AP-1) (**Figure 3F**), which plays a role in VEGF-mediated endothelial cell migration and proliferation[51]. *Socs 3 (*suppressor of cytokine signaling) was also increased in the PND 1 female gCaps (**Figure 3F**) and it decreases inflammation-mediated pathological angiogenesis [52–55]. Endothelial *Socs 3* limited inflammation and promoted survival in endotoxin-exposed mice [56]. MHC-II genes such as *H2-Aa* and *H2-Ab1* were exclusive to the male gCaps at PND 21 (**Figure 3H**). Endothelial expression of MHC-II genes may point to their role as antigen-presenting cells to T-cells and furthers the endothelial to immune cell link [57–60], with interferon-gamma mediated signaling inducing the expression of MHC class II genes. Finally, differentially expressed genes in arterial cells between the male and female lung at PND 21 along with enriched biological pathways are shown in **Supplemental Figure 2**.

### Sex specific transcriptional signature in lung immune cells during development

Next, we sub clustered the lung immune cells and identified 12 and 14 distinct populations at PND 1 (**Figure 4A**) and PND 21 (**Figure 4B**) respectively. At PND 1, we identified a population of transient *Galanin +* (*Gal*) and *Spint1 +* macrophages (**Figure 4C**) that were not present at the subsequent time-points. This population of macrophages has been reported previously by Domingo-Gonzalez et al. [33] as being present in the lung at a similar developmental stage around small pulmonary vessels. We also identified high expression of *Angptl4* (Angiopoietin-like 4) in this cluster (**Figure 4C, right**), which is a potent angiogenic factor [61] and has anti-inflammatory activity [62,63]. Sex-specific differences in transient macrophages identified biological pathways related to TNF-alpha signaling and response to cytokines, that were enriched in female cells (**Figure 4D, Supplemental Table 5**).

**Figure 4:**
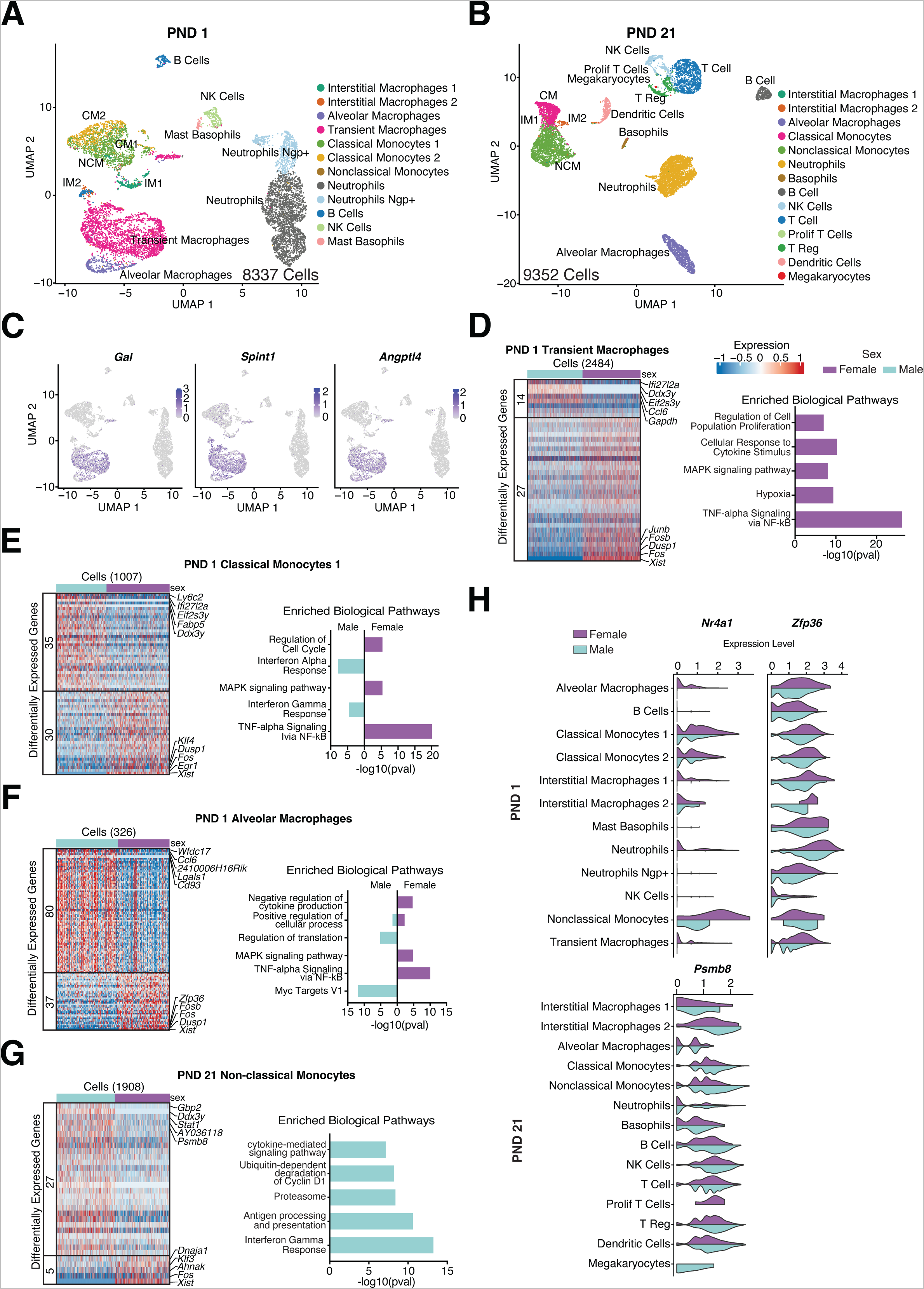
Sex-specific programming of biological pathways in the lung immune cells during lung development. **(A-B)** UMAP of sequenced lung immune cells at PND 1 and PND 21. **(C)** Feature plots showing high expression of *Gal, Spint 1*and *Angptl4* in the transient macrophage subcluster. **(D)** Differential expression between male and female cells and enriched biological pathways in transient macrophages. **(E)**. Heatmap of differentially expressed genes between male and female cells with enriched biological pathways in PND 1 classical monocytes 1, **(F)** PND 1 alveolar macrophages and **(G)** PND 21 non-classical monocytes. **(H)** Violin plots of differentially expressed genes in the male and female lung immune cells *Nr4a1, Zfp36* and *Psmb8*. Number of cells sequenced are shown within parentheses.

We also identified two distinct clusters of classical monocytes (CM) in the PND 1 lung, that we labeled CM1 and CM2. CM1 was the traditional classical monocyte population. The CM2 cluster however, expressed several markers like *Elane, Prtn3, Lcn and Fcnb*, which were recently described in a population of “neutrophil-like monocytes” derived from granulocyte-monocyte progenitors [64,65] and are proposed to have immunoregulatory functions [66]. Differentially expressed genes and differentially modulated biological pathways in PND 1 classical monocytes 1 (**Supplemental Table 6**), PND 1 alveolar macrophages (**Supplemental Table 7**), and PND 21 non-classical monocytes (**Supplemental Table 8**) were identified **(Figure 4E-G)**. Genes classified as immediate early response genes (IEGs); *Fos, Fosb, Jun,* and *Egr1* were enriched in the female immune cells. IEGs are a group of genes that are rapidly and transiently activated in response to various stimuli, including growth factors, stress, and other extracellular signals. They are called “immediate early” because their expression can be detected within minutes to a few hours after the stimulus [67]. IEGs play a crucial role in initiating and coordinating cellular responses to environmental changes. They act as primary response genes and are often involved in regulating the expression of downstream target genes, including other transcription factors, signaling molecules, and structural proteins. Their activation serves as an early and rapid signaling event that triggers a cascade of molecular and cellular responses [67]. Sex-specific differences in IEGs have been reported in the brain [68–71]. *Egr1* plays a fundamental role in monocytic commitment and macrophage polarization [72,73]. Trizzino *et al* showed that enhancers associated with the inflammatory response are EGR1 target regions in macrophages, and *Egr1* represses inflammatory enhancers in developing and mature macrophages, thus attenuating the immune response [73].

A common overarching finding across all these cell-types was the role of TNF-alpha signaling pathway in female immune cells and the interferon pathway in the male immune cells. **Figure 4H** shows the expression levels of genes *Nr4a1* or *Nur77* involved in the TNF-alpha signaling pathway that regulates the expression of IEGs [74] and plays a role in macrophage polarization [75] with female lung immune cells showing higher *Nr4a1* expression. Anti-inflammatory and anti-oxidant role of *Nr4a1* have been reported in previous studies[76,77]. RNA-binding protein tristetraprolin (TTP, encoded by the *Zfp36* gene) is expressed higher in female cells (**Figure 4H**) and inhibits TNF-alpha mediated cytotoxicity in immune cells [78,79]. *Psmb8* (**Figure 4H**), also known as β5i or LMP7, is a specialized subunit of the immunoproteasome, which is a variant of the proteasome involved in antigen processing and presentation in immune cells and is reported to play role in chronic lung diseases [80]. It is induced by IFN-gamma in lung immune cells [81,82]. *Psmb8* had a greater expression in male IMs, classical and nonc-lassical monocytes at PND21, compared to female. Differentially expressed genes and biological pathways in male and female neutrophils and T cells are included in **Supplemental Figure 3 A-B** (**Supplemental Table 9-10**).

### Neonatal hyperoxia exposure during the saccular stage of lung development leads to persistent sex-specific changes in the transcriptional state of cell sub-populations

To discern the effects of neonatal hyperoxia exposure during the saccular stage of lung development, we performed sc-RNASeq from room air and hyperoxia-exposed male and female murine lungs at PND 21 (**Figure 5A**). UMAP plots of the lung cell sub-populations in room air and hyperoxia are shown in **Figure 5B**. The number of differentially expressed genes (DEGs) between room air and hyperoxia-exposed mice (both sexes combined) at PND 21 in different endothelial and immune cells is shown in **Figure 5C** (**Supplemental Table 11**). gCaps, alveolar macrophages, non-classical monocytes and neutrophils show the greatest number of DEGs among the highlighted cell subpopulations. To link the changes in gene expression by biological sex immediately after hyperoxia exposure at PND 7 and after the organ has been allowed to recover in room air at PND 21, we compared the DEGs in endothelial and immune cells that are sex-specific (exclusive to male or female lung) or common (common to male and female lung) in **Figure 5D**. The sex-specific differences in gene expression in the hyperoxia exposed lung was particularly striking among the lung immune cells. Alveolar macrophages and T cells are predominated by a female response, while B-cells and classical monocytes were male biased in gene expression changes at PND21.

**Figure 5:**
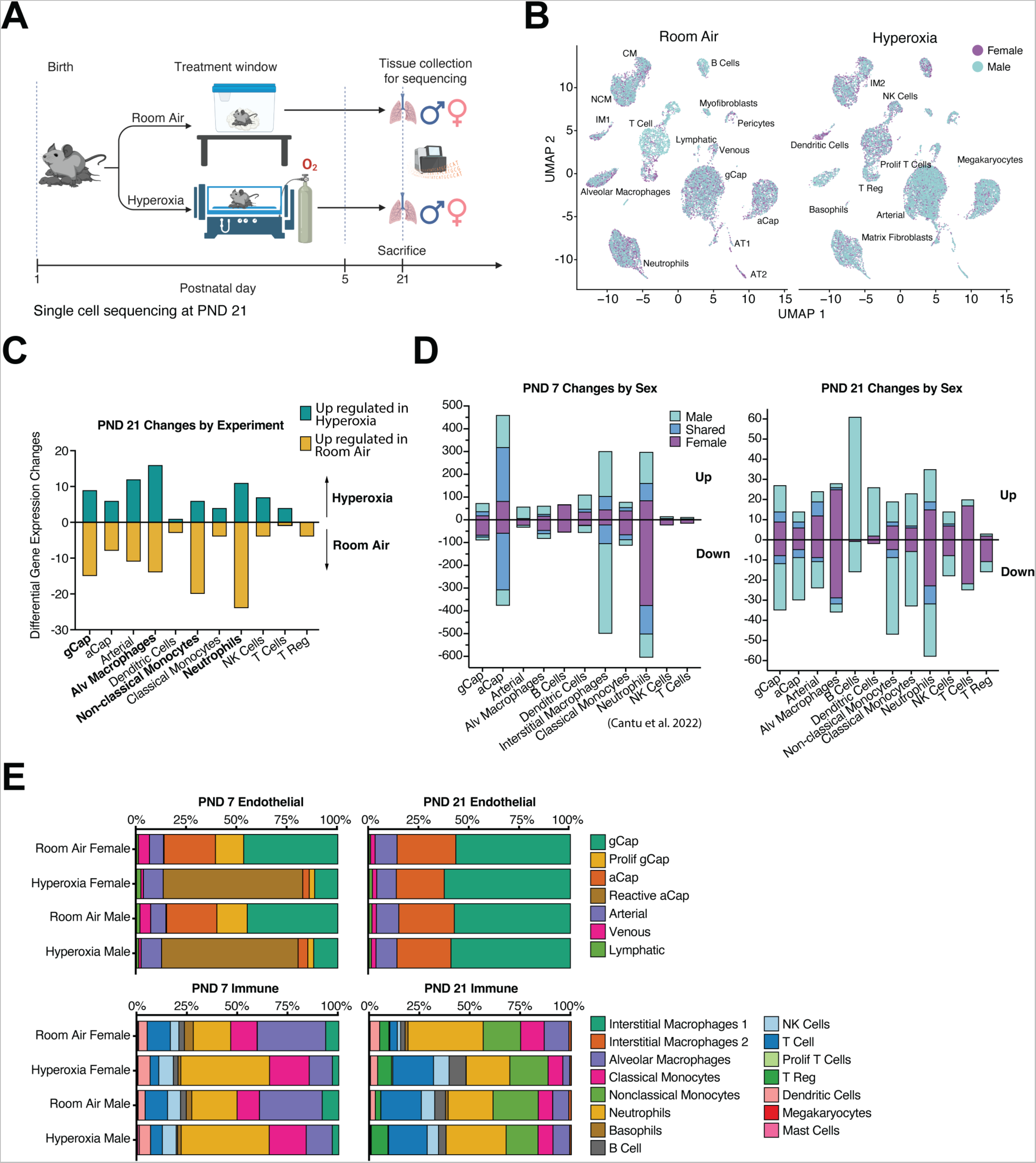
Neonatal hyperoxia exposure during the saccular stage of lung development leads to persistent sex-specific changes in the transcriptional state of lung cells. **(A)** Experimental design and timeline. Mouse pups (C57BL6) from multiple litters were pooled before being randomly and equally redistributed to two groups, one group exposed to room air (21% O2) and the other group exposed to hyperoxia (95% O2), within 12 h of birth for 5 days, and euthanized on PND 21. **(B)** UMAP of sequenced lung cells at PND21 identified distinct cell clusters split by sex and experimental condition (room air and hyperoxia). **(C)** Number of differentially expressed genes (DEGs) among endothelial and immune cells (upregulated on top and downregulated at the bottom) in response to hyperoxia in both male and female mice (sexes combined). **(D)** DEGs between hyperoxia and room air that are either shared between male and female (dark blue), unique in male (pale blue) or unique in female (purple) in lung endothelial and immune cells at PND 7 and PND 21 **(E)** The relative changes in lung endothelial and immune cells at PND 7 (immediately after hyperoxia exposure) and PND 21 (after recovery in room air).

Remarkable changes in cellular composition of the lung after hyperoxia exposure have been reported in prior studies [23,35,83,84]. The relative changes in lung endothelial and immune cell number at PND 7 (immediately after hyperoxia exposure) and PND 21 (after recovery in room air) are shown in **Figure 5E**. gCaps were decreased, while reactive aCaps increased significantly at PND 7. However, at PND 21, the endothelial cells relative composition was similar between the room-air and hyperoxia-exposed lung in male and female lung. The lung immune cells showed increase in neutrophils and a decrease in alveolar macrophages and T cells at PND 7. Interestingly, at PND 21 the female hyperoxia-exposed lung shows increased T-cells and B-cells and decreased alveolar macrophages and neutrophils compared to males. These results must be considered speculative as validation of these findings with cell-specific markers was not performed in this study. However, this highlights that while the endothelial cells may recover in terms of relative composition by PND 21, persistent changes are seen among the lung immune cells.

### Alterations in gene expression in lung endothelial cells in response to neonatal hyperoxia exhibit sex-specific differences

UMAP plot of different lung endothelial cell populations (**Figure 6A**) at PND 21 and split by treatment (room air and hyperoxia) and sex (male and female) are shown in **Figure 6B**. Differentially expressed genes (DEGs), overlap or lack thereof between male and female lung endothelial cells and enriched biological pathways in the male and female hyperoxia- exposed lung gCaps, aCaps and arterial endothelial cells compared to room air controls are shown in **Figure 6C-E** (**Supplemental Figure 4, Supplemental Table 12-14**). The number of differentially expressed genes are less compared to our previously published results at PND 7 (immediately after hyperoxia exposure)[23]. Venn diagrams (**Figures 6C-E, center**) show greater differences than similarities between male and female lung endothelial cells with majority of the DEGs being autosomal (not present on the sex chromosomes). One common finding across all endothelial cells was the upregulation of *Cdkn1a* or *p21*. Induction of *p21* in the neonatal lung following hyperoxia exposure has been reported before and is protective against oxygen-induced toxicity [85–87]. Expression of the long non-coding rna *Xist (*X-inactive specific transcript) was decreased in the female endothelial cells upon exposure to hyperoxia (**Figure 6F**). *Xist* is responsible for initiating and maintaining the inactivation of one X chromosomes in female cells. Decreased *Xist* expression attenuates angiogensis [88–90]. Interestingly, *Xist* expression is induced by glucocorticoids [91]. Hyperoxia-exposed gCaps showed increased expression of insulin like growth factor binding protein 7 (*Igfbp7),* also known as angiomodulin (**Figure 6C**). *Igfbp7* is expressed in the developing vasculature and plays a role in immune cell extravasation in disease states[92,93]. *Hspb1* (Heat Shock Protein Family B member 1) was upregulated in hyperoxia-exposed aCaps (**Figure 6C**) and has been shown to regulate VEGF-mediated angiogenesis and inhibit endothelial-to-mesenchymal transition [94,95]. Pathways related to interferon gamma and antigen presentation were downregulated in the male hyperoxia-exposed endothelial cells (**Supplemental Figure 4**), while the TNF-alpha signaling via NF-kappaB was positively modulated in the male hyperoxia-exposed gCaps (**Figure 6C**). Interestingly, this was opposite to what was observed in room air male lung endothelial cells, described in the prior sections.

**Figure 6:**
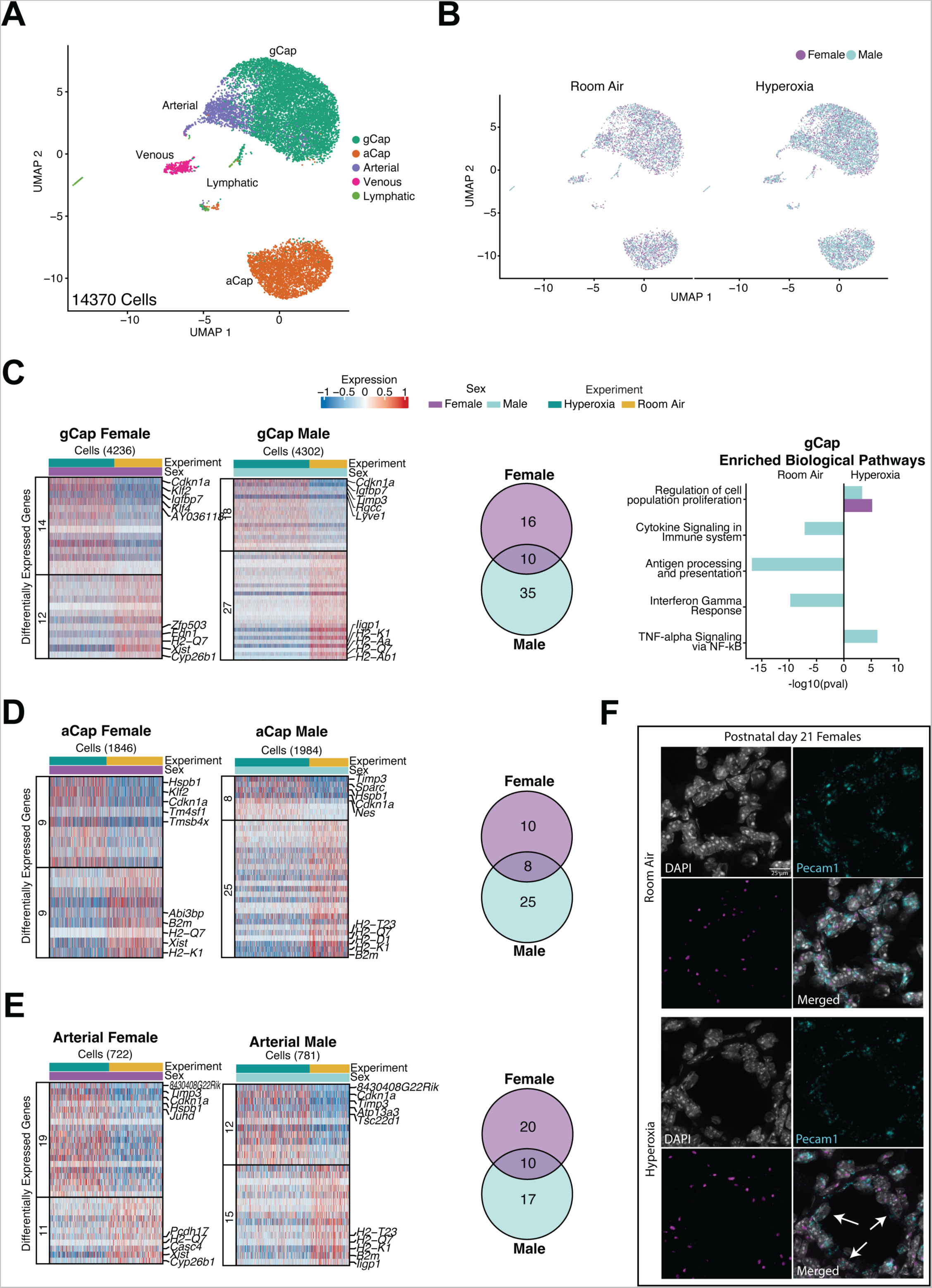
Sex-specific changes in the lung endothelial cells at PND21 in response to neonatal hyperoxia during the saccular stage of lung development. **(A)** UMAP of sequenced lung endothelial cells identified five distinct clusters at PND 21. **(B)** UMAP of lung endothelial cells split by treatment (room air and hyperoxia) and sex at PND 21. **(C)** Heatmap of differentially expressed genes (room air vs. hyperoxia) in female and male gCaps, Venn diagram showing the number of unique (sex-specific) and common DEGs and enriched biological pathways. **(D)** Heatmap of differentially expressed genes (room air vs. hyperoxia) in female and male aCaps and Venn diagram showing the number of unique (sex-specific) and common DEGs. **(E)** Heatmap of differentially expressed genes (room air vs. hyperoxia) in female and male arterial endothelial cells and Venn diagram showing the number of unique (sex-specific) and common DEGs. Number of cells sequenced are shown within parentheses. **(F)** *In situ* hybridization of *Pecam1* and *Xist* in room air and hyperoxia PND 21 female lungs. Arrows pointing to *Pecam1*+ cells that show little or no *Xist* signal.

### Sex-specific changes in lung immune cells in response to neonatal hyperoxia

UMAP plot of different lung immune cell populations (**Figure 7A**) at PND 21 split by treatment (room air and hyperoxia) and sex (male and female) are shown in **Figure 7B**. We identified two populations of interstitial macrophages in this study (IM1 and IM2). Heatmap showing differential expression of genes between these two IM clusters is shown in **Supplemental Figure 5A** (**Supplemental Table 15**). Interestingly, these correspond to CD206+ (IM1, higher expression for *Mrc1*, *CD163*) and CD206- (IM2, higher expression of *MHCII* markers, *CD74*) interstitial macrophages reported in the murine lung with distinct biological niches and functional properties[96]. Also, our previous study that assessed the changes in the hyperoxia-exposed lung immediately after hyperoxia exposure at PND 7 identified a population of alveolar macrophages that had higher expression of *Inhba* [23]. This was also reported by other studies[35]. **Figure 7C** shows the changes in the relative contribution of AMs and *Inhba+* AMs across three developmental time-points with and without hyperoxia exposure. AMs are depleted in the lung after hyperoxia exposure but recover by PND 21. *Inhba+* AMs increase in the hyperoxia-exposed lung at PND 7 but returns to baseline by PND 21. IM1s decrease progressively through PND 1-21, while relative contribution of IM2s increases. Hyperoxia exposure attenuates the decrease in IM1 and increase in IM2 at PND 7 (**Figure 7D**). As described previously, the two identified classical monocytes clusters at PND 1; CM1 and CM2 (“neutrophil-like”) [64] follow increasing and decreasing trends in relative abundance across the three time-points in early lung development and are not perturbed by hyperoxia exposure (**Figure 7E**).

**Figure 7:**
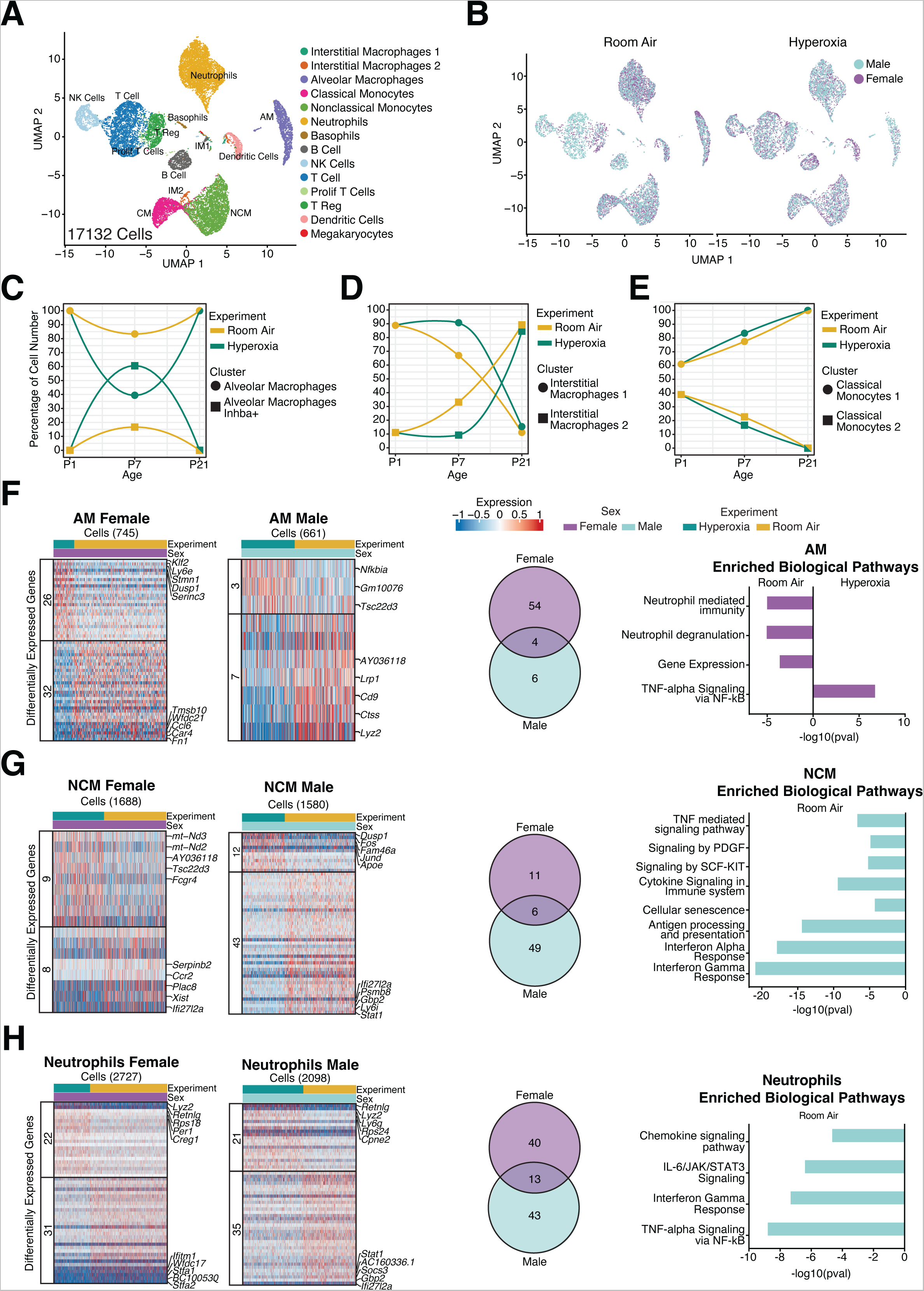
Sex-specific changes in the immune cells at PND 21 in response to neonatal hyperoxia during the saccular stage of lung development. **(A)** UMAP of sequenced lung immune cells identified 14 distinct clusters at PND 21. **(B)** UMAP of lung immune cells split by treatment room air and hyperoxia and sex at PND 21. **(C)** Relative variation in cell composition in room air or hyperoxia of alveolar macrophages (AMs) and *Inhba*+ AMs, **(D)** interstitial macrophage 1 (IM1) and IM2, **(E)** classical monocyte 1 (CM1) and CM2 at PND 1, PND 7 and PND 21. **(F)** Heatmap showing differentially expressed genes between room air and hyperoxia in female and male, Venn diagram showing the number of unique (sex-specific) and common DEGs in hyperoxia-exposed lungs and enriched biological pathways in alveolar macrophages, **(G)** non-classical monocytes and **(H)** neutrophils. Number of cells sequenced are shown within parentheses.

Differentially expressed gene (DEGs) overlap or lack thereof between male and female lung immune cells and enriched biological pathways in hyperoxia-exposed lung alveolar macrophages (**Figure 7F, Supplemental Table 16**), non-classical monocytes (**Figure 7G, Supplemental Table 17**), neutrophils (**Figure 7H, Supplemental Table 18**) and classical monocytes (**Supplemental Figure 5B, Supplemental Table 19**) were determined. The number of differentially expressed genes is lower when compared to our previously published findings at PND 7[23]. Additionally, the Venn diagrams highlight a greater number of distinctions rather than similarities between male and female lung immune cells with most DEGs being autosomal (not present on the sex chromosomes). Biological pathways in female alveolar macrophages included TNF-alpha signaling via NF-kappa B that was positively modulated and neutrophil mediated immune pathways that were negatively enriched (**Figure 7F, right**). Interferon mediated pathways and antigen processing and presentation were negatively modulated in the male non-classical monocytes (**Figure 7G, right**). Expression of *Xist* was decreased in the female classical (**Supplemental Figure 5B**) and non-classical monocytes (**Figure 7G**). Expression from the inactive X chromosome in immune cells increases the female predisposition to autoimmune conditions[97], but the epigenetic features of the inactive X-chromosome varies in different classes of immune cells[98]. Both pro- and anti-inflammatory roles of *Xist* have been reported in the literature [99–102]. Nevertheless, the modulation of *Xist* expression in female lung immune cells by hyperoxia exposure points to its probable role in the biological response.

On the other hand, expression of another X-chromosome gene, Tsc22 domain family protein 3 (*Tsc22d3*; also known as glucocorticoid-induced leucine zipper, *Gilz*), was increased in female lung alveolar macrophages and non-classical monocytes in the female lung (**Supplemental Figure 5C**). We validated the expression of *Tsc22d3* room air murine lung and showed expression in *Adgre1*+ lung macrophages and *Siglecf*+ alveolar macrophages (**Supplemental Figure 5D-E**). Interestingly, *Tsc22d3* is a glucocorticoid-inducible transcriptional regulator, and functions as a potent mediator of anti-inflammation [103–106].

### Pseudotime analysis reveals transcriptional changes in lung capillary endothelial cells through saccular and alveolar stages of development

To investigate common molecular mechanisms that guide the development and maturation of lung capillary endothelial cells, we next examined transcriptional changes in this population throughout their developmental progression. We used Monocle3 to identify and order developing lung capillary endothelial cells in ‘‘pseudotime’’. Monocle3 computationally orders cells in an unsupervised manner by maximizing the transcriptional similarity between successive pairs of cells. **Supplemental Figure 6A** shows the UMAP of lung aCaps and gCaps from PND 1 through PND 21. It was evident that the aCaps and gCaps clustered closer during the earlier time points but became transcriptionally distinct by PND 21. For further analysis, we kept the aCaps and gCaps at PND1 but focused on the aCaps at subsequent time-points since our previous study found that aCaps showed the greatest number of DEGs at PND 7 after hyperoxia exposure [23]. Lung capillary endothelial cells were ordered along a putative developmental trajectory (pseudotime) from least to most differentiated (**Figure 8A**). Interestingly, the endothelial cells fit the developmental timeline with the cells at PND1 being the least differentiated and the cells at PND21 being further along the pseudotime trajectory, suggesting that distinct molecular pathways guide the maturation of these endothelial cells. Next, we analyzed the genes involved in cell maturation that deviated under hyperoxia conditions. The Pseudotime trajectory of these cells is shown in **Figure 8B**. Though the Pseudotime values fit with the developmental timeline, the pattern looks different from the room air endothelial cells, with a bifurcation from PND1 to PND7 and PND21 rather than a sequential trajectory in room air cells (**Figure 8A**). To determine which genes, regulate the maturation and recovery after injury of lung aCaPs, we performed hierarchical clustering of the genes whose expression varied as a function of pseudotime. We then subdivided these genes into those that were exclusive to room air or hyperoxia exposed lungs (**Figure 8C**).

**Figure 8:**
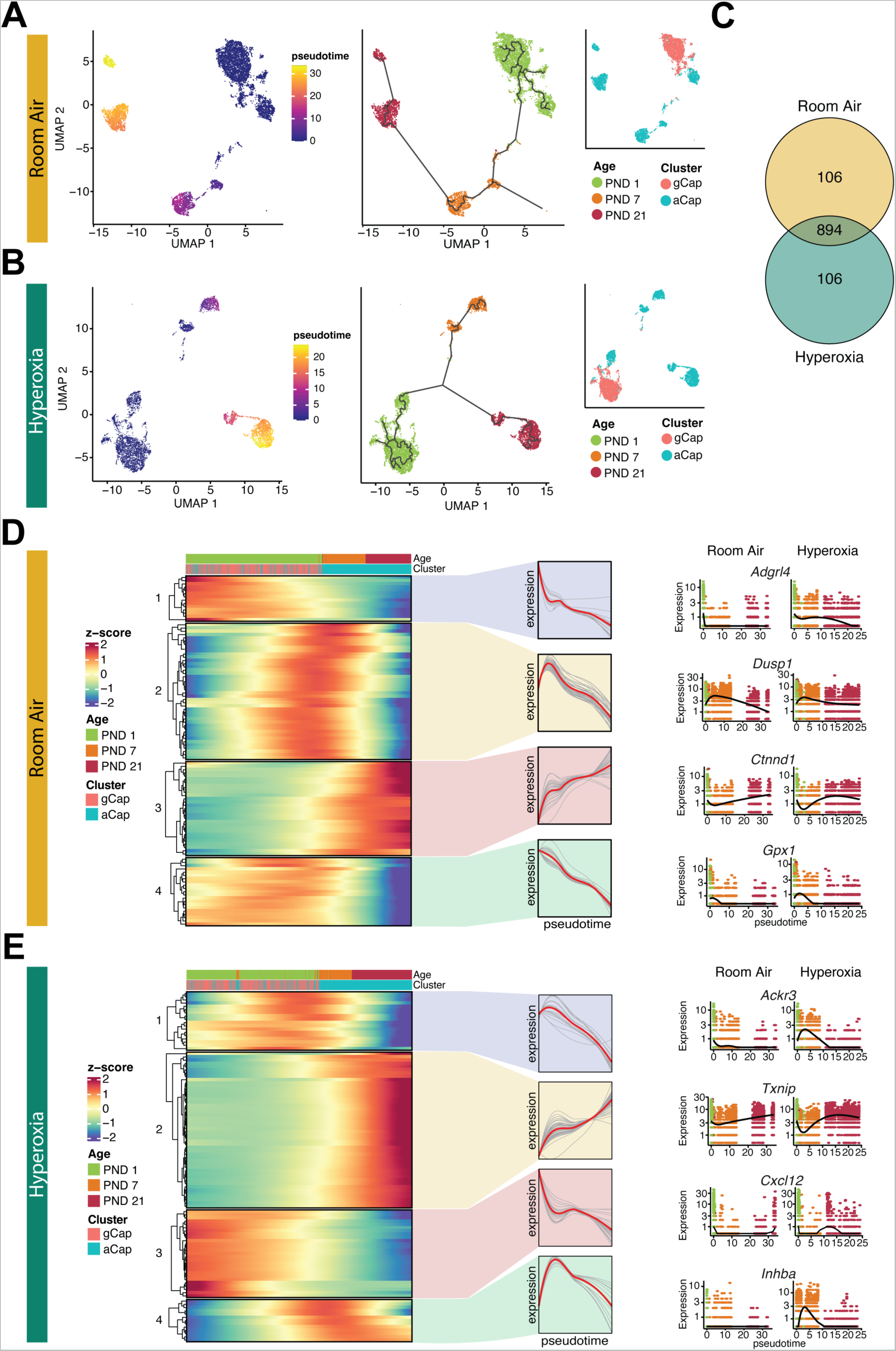
Pseudotime analysis reveals transcriptional changes in lung capillary endothelial cells exposed to hyperoxia throughout development. **(A)** Monocle3 pseudotime trajectory of pulmonary capillary endothelial cells in room air-exposed lungs from PND 1 through PND 21. Cells are colored by pseudotime score, with dark colors representing immature cell stages and light colors representing mature cell stages (left) along with age (middle) and cluster name (right). **(B)** Monocle3 pseudotime trajectory of pulmonary capillary endothelial cells in hyperoxia-exposed lungs (PND1-room air, PND 7-hyperoxia and PND 21-hyperoxia) from PND 1 through PND 21. Cells are colored by pseudotime score, with dark colors representing immature cell stages and light colors representing mature cell stages along with age (middle) and cluster name (right). **(C)** Venn diagram of top 1000 genes that change as a function of pseudotime in room air (top) and hyperoxia (bottom). **(D)** Distinct clusters of pseudotime-dependent genes (exclusive to room air) with dynamic expression patterns plotted across pseudotime as heatmaps, with blue indicating low levels and red indicating high levels of expression (left). Gene expression trends for each gene shown in black with the average expression line highlighted in red (middle). Differential expression patterns of one example gene from each group of genes (right). **(E)** Distinct clusters of pseudotime-dependent genes (exclusive to hyperoxia) with dynamic expression patterns plotted across pseudotime as heatmaps, with blue indicating low levels and red indicating high levels of expression (left). Gene expression trends for each gene shown in black with the average expression line highlighted in red (middle). Differential expression patterns of one example gene from each group of genes (right).

The heatmap in **Figure 8D-E** shows the distinct clusters of pseudotime-dependent genes and their expression trends. The expression pattern of one example gene from each cluster is shown along with its relative expression both in room air and hyperoxia (**Figure 8D-E, right**). *Adgrl4* (Adhesion G Protein-Coupled Receptor L4), also known as the endothelial orphan receptor ELTD1, plays an important pro-angiogenic role [107]. The expression is high at PND 1 and decreases significantly in room air through PND21. In the hyperoxia-exposed endothelial cells, the expression is sustained through the early alveolar stages of lung development. *Dusp1* or *Mkp-1* expression, which also is pro-angiogenic [108], was also affected similarly with a sustained higher expression in hyperoxia exposed endothelial cells compared to room air controls. CTNND1 encodes a cellular adhesion protein p120-catenin (p120) which is essential for vascularization and activation of Wnt signaling activity by decreasing β-catenin degradation [109]. Interestingly, the expression pattern of this gene is lower at PND 21 in the hyperoxia-exposed lung compared to room air controls in which the expression levels continue to increase from PND 1 through PND 21. Lastly, glutathione peroxidase-1 (*Gpx1*), with an antioxidant role [110,111], was increased to a greater extent at PND 1 and decreased continuously through PND 21 under room air conditions. In hyperoxia-exposed endothelial cells, the expression was increased to a greater extent after hyperoxia exposure but returned to baseline at PND 21. In hyperoxia (**Figure 8E**), *Ackr3* or *CXCR7*, receptor for the pro-angiogenic ligand CXCL12 [112,113] was increased in the hyperoxia exposed endothelial cells immediately after hyperoxia compared to room air controls where the levels were low. Compared to room air endothelial cells, expression of thioredoxin-interacting protein *Txnip* was decreased immediately after hyperoxia exposure followed by a brief increase in expression in hyperoxia-exposed endothelial cells. Endothelial *Txnip* inhibits Hif-1 alpha and causes endothelial dysfunction [114–116]. *Cxcl12*, a chemokine with pro-angiogenic effects [117,118], was transiently increased in the hyperoxia-exposed lungs but stayed consistently low in the room air group. Expression of *Inhba* was increased transiently after hyperoxia exposure returned to baseline at PND 21. We and others have reported on increased expression of *Inhba* in the capillary endothelium [23,35], which has an anti-angiogenic role [119].

### Pseudotime analysis reveals transcriptional changes in lung alveolar macrophages throughout development and perturbations by early hyperoxia exposure

Using Monocle 3, we examined transcriptional changes in lung alveolar macrophages in the postnatal lung saccular, early alveolar, and late alveolar stages of development. We included the population of transient macrophages identified at PND 1 and analyzed the cells from room air controls (**Figure 9A**) and hyperoxia exposed lungs (**Figure 9B**). Lung alveolar macrophages were then ordered along a pseudotime trajectory from least to most differentiated. Interestingly, the transient and alveolar macrophages fit the developmental timeline (transient macrophages to alveolar macrophages) with the cells at PND 1 being the least differentiated and the cells at PND 21 being further along the trajectory under both room air- and hyperoxic conditions, suggesting that distinct molecular pathways guide the maturation of these cells. To determine which genes, regulate the maturation and recovery after injury of lung alveolar macrophages we performed hierarchical clustering of the genes whose expression varied as a function of pseudotime. We then subdivided these genes into those that were exclusive to room air or hyperoxia exposed lungs (**Figure 9C**).

**Figure 9:**
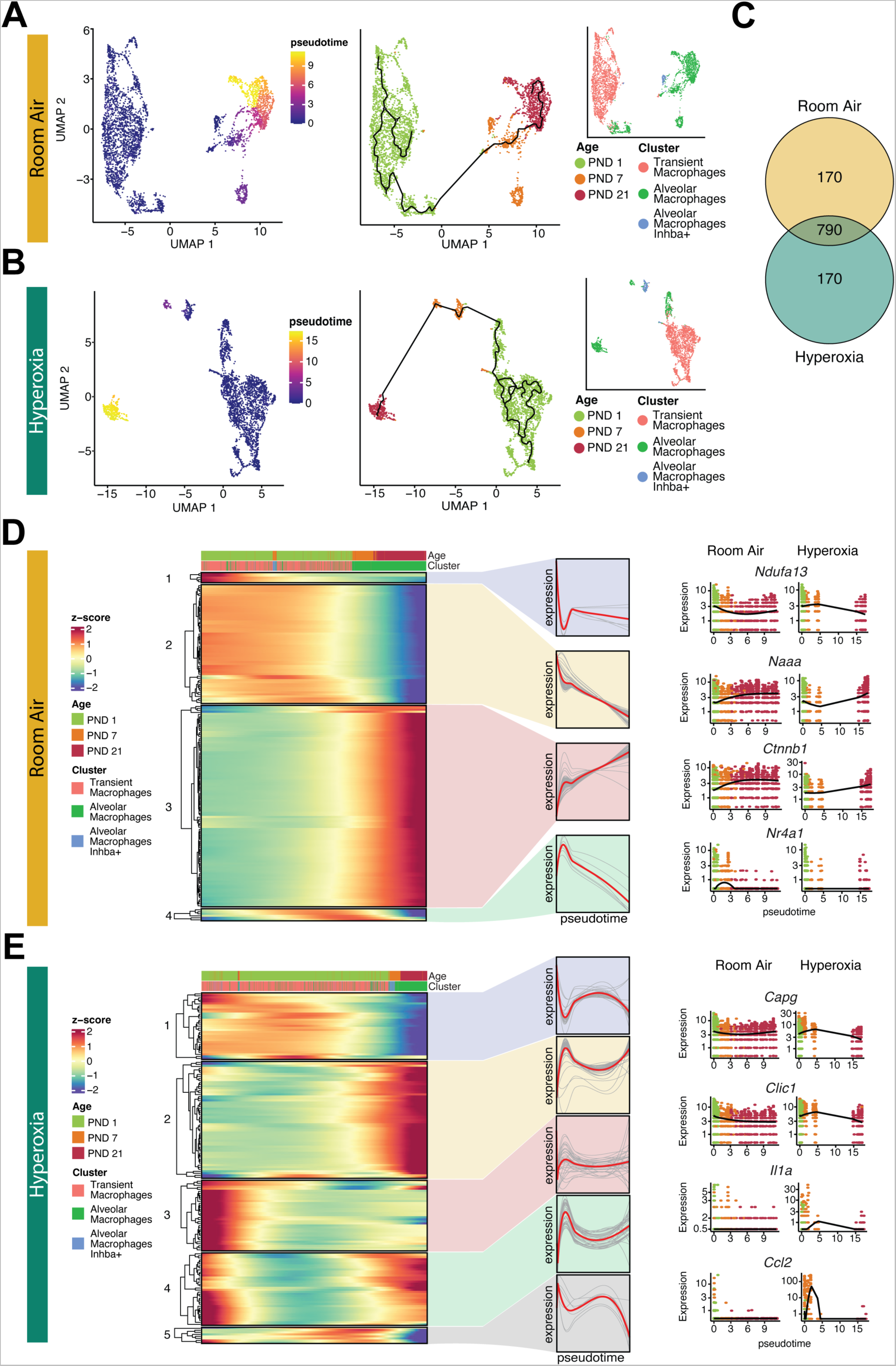
Pseudotime analysis reveals transcriptional changes in lung transient and alveolar macrophages through saccular and alveolar stages of lung development. **(A)** Monocle3 pseudotime trajectory of pulmonary transient and alveolar macrophages in room air-exposed lungs from PND 1 through PND 21. Cells are colored by pseudotime score, with dark colors representing immature cell stages and light colors representing mature cell stages (left) along with age (middle) and cluster name (right). **(B)** Monocle3 pseudotime trajectory of pulmonary alveolar macrophages in hyperoxia-exposed lungs (PND 1-room air, PND 7-hyperoxia and PND 21-hyperoxia) from PND 1 through PND 21. Cells are colored by pseudotime score, with dark colors representing immature cell stages and light colors representing mature cell stages (left) along with age (middle) and cluster name (right). **(C)** Venn diagram of top 1000 genes that change as a function of pseudotime in room air (top) and hyperoxia (bottom). **(D)** Distinct clusters of pseudotime-dependent genes (exclusive to room air) with dynamic expression patterns plotted across pseudotime as heatmaps, with blue indicating low levels and red indicating high levels of expression (left). Gene expression trends for each gene shown in black with the average expression line highlighted in red (middle). Differential expression patterns of one example gene from each group of genes (right). **(E)** Distinct clusters of pseudotime-dependent genes (exclusive to hyperoxia) with dynamic expression patterns plotted across pseudotime as heatmaps, with blue indicating low levels and red indicating high levels of expression (left). Gene expression trends for each gene shown in black with the average expression line highlighted in red (middle). Differential expression patterns of one example gene from each group of genes (right).

The heatmap in **Figure 9D** shows distinct clusters of pseudotime-dependent genes exclusive to room air and their gene expression trends. The expression pattern of one example gene from each cluster is shown with its relative expression in both room air and hyperoxia-exposed lung alveolar macrophages. NDUFA13 (also known as GRIM-19), is an essential subunit of mitochondrial respiratory chain complex I, essential for IL-1β production, and generation of mitochondrial reactive oxygen species[120]. This gene had a decreasing expression pattern across the three time-points in room air. However, in hyperoxia, there is an upregulation at PND 7 with a return to room air levels at PND 21. N-acylethanolamine acid amidase (*Naaa*) that plays a role in macrophage activation [121,122], shows a pattern of sustained increase expression in room air. Post hyperoxia-exposure, the levels increase at PND 7 and a return to room air levels by PND 21. Beta-catenin (*Ctnnb1*) plays a role in macrophage lineage differentiation [123] and myeloid β-catenin expression modulates macrophage inflammatory responses[124]. This gene had reduced expression at PND 7 in hyperoxia exposed cells but returned to room air levels by PND 21. *Nr4a1* (or *Nur77*) is a key transcriptional regulator and induces an anti-inflammatory metabolic state in macrophages [125]. Expression was initially low, rising at PND 7 and decreasing by PND 21 in room air lungs, this pattern was not observed in hyperoxia exposed lungs. Hierarchical clustering of the genes exclusive to hyperoxia whose expression varied as a function of pseudotime, and their expression trends is shown in **Figure 8D**. The macrophage capping protein *CapG* belongs to the gelsolin superfamily and is a redox-sensitive protein [126]. This gene showed an expression pattern of high baseline levels at PND 1 and decreased expression at PND 21 after hyperoxia exposure. Intracellular chloride channel protein 1 (*Clic1*) showed a similar pattern and plays a role macrophage reactive oxygen species production and is pro-inflammatory[127–129]. IL-1 alpha, which contributes to neutrophil recruitment in the hyperoxia-exposed lung [130,131], showed increased expression at PND 7 with a return to room air levels by PND 21. *CCl2* or monocyte chemoattractant protein-1, MCP-1 was also increased at PND 7 in the hyperoxia-exposed lung with return to baseline by PND 21.

### Novel endothelial-immune interactions in the saccular stage of lung development

We used the CellChat bioinformatic package to infer, visualize and analyze intercellular communications from scRNA-seq data. We interrogated the cell-cell interactions between the endothelial and immune cells at PND 1 room air and PND 21 room air and hyperoxia cells and sought to identify pathways that were sex-specific. The cell-cell communication between the different endothelial and immune cell subpopulations at PND 1 is summarized in **Figure 10**. We identified the global communication patterns that connect cell groups with signaling pathways in the context of outgoing or incoming signaling. (**Supplemental Figure 7A-B**). *<u>The outgoing signaling shows six patterns.</u>* The endothelial cells separated into two main patterns: with the aCaPs in pattern 1, and gCaPs/arterial/venous/proliferating ECs in pattern 5. Among the immune cells, neutrophils (pattern 2), transient/alveolar macrophages (pattern 3), interstitial macrophages (pattern 4), and B/NK cells (pattern 6) formed their respective outgoing patterns. The signaling molecules associated with these patterns are highlighted in **Supplemental Figure 7A**. *<u>The incoming signaling patterns</u>* grouped the endothelial cells; gCaP/proliferating ECs/arterial (pattern #1), venous (pattern #6) and the aCaPs (pattern #4) in three groups. Macrophages (pattern #3), monocytes and neutrophils (pattern #2), and NK cells (pattern#5), comprised the three patterns among immune cells (**Supplemental Figure 7B**). The signaling molecules associated with these patterns are highlighted in **Supplemental Figure 7B**. The number and strength of interactions among and between endothelial and immune cells is shown in **Figure 10A**. The lung endothelial and immune cells show robust communication patterns with autocrine signaling among themselves. The alveolar and transient macrophages have a high interaction number with endothelial cells, while the classical monocytes and neutrophils display a higher interaction weight/strength, pointing to the role of these intercellular communications patterns during postnatal lung development.

**Figure 10:**
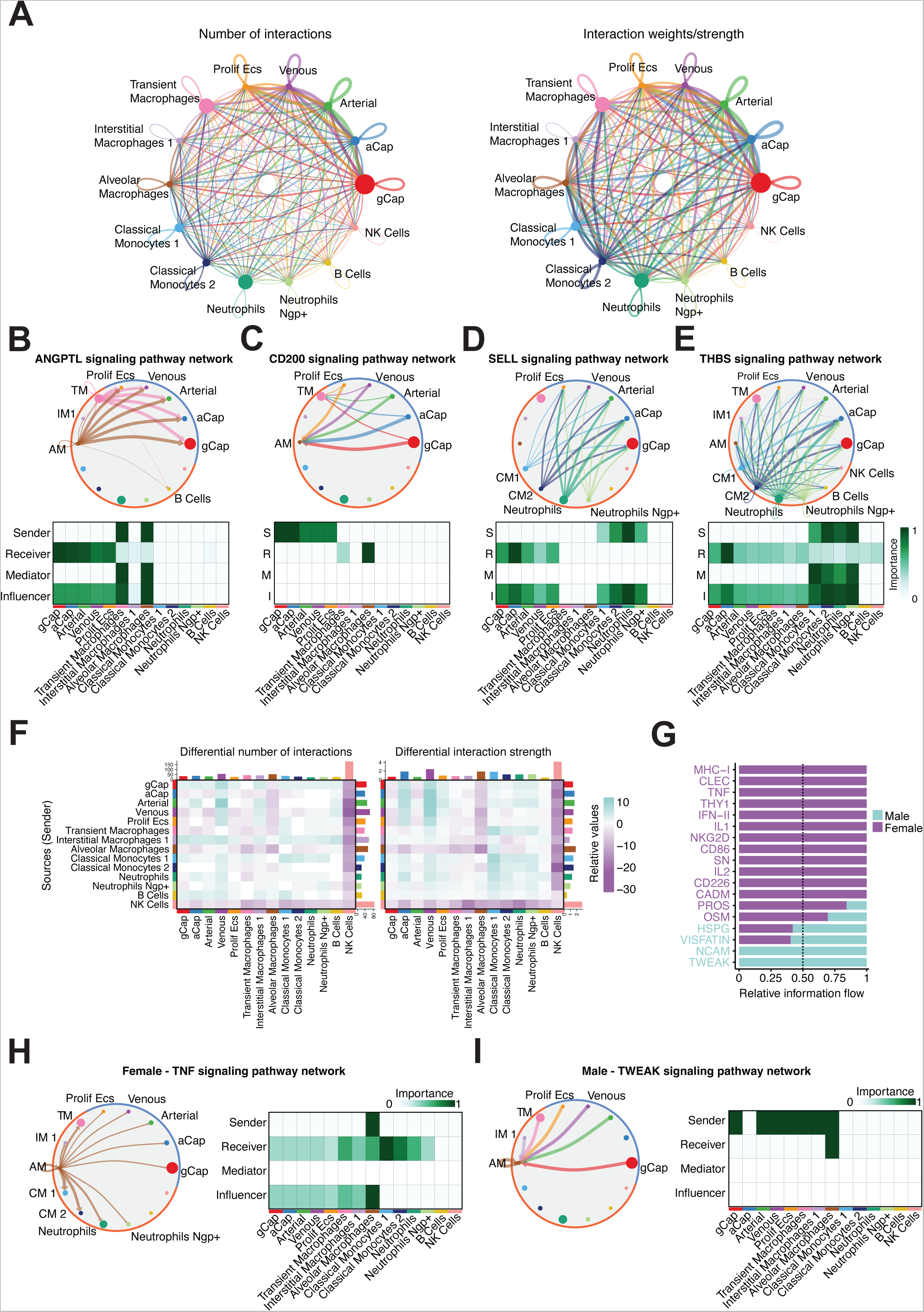
Novel endothelial-immune interactions in the postnatal lung at PND 1. **(A)** Circle plot showing the number of interactions between endothelial and immune cells. Edge colors are consistent with the sources as sender, and edge weights are proportional to the interaction strength. Thicker edge line indicates a stronger signal. Circle sizes are proportional to the number of cells in each cell group. **(B)** Circle plot shows the inferred intercellular communication network for *Angptl* signaling highlighting the paracrine signaling to endothelial cells (top). Circle sizes are proportional to the number of cells in each cell group and edge width represents the communication probability. Edge colors are consistent with the signaling source. Heatmap shows the relative importance of each cell group based on the computed four network centrality measures of *Angptl* signaling network (bottom). **(C)** Circle plot shows the inferred intercellular communication network for *Cd200* signaling highlighting the paracrine signaling to immune cells (top). Heatmap shows the relative importance of each cell group based on the computed four network centrality measures of *Cd200* signaling network (bottom). **(D)** Circle plot shows the inferred intercellular communication network for *SELL* signaling highlighting the paracrine signaling to endothelial cells (top). Heatmap shows the relative importance of each cell group based on the computed four network centrality measures of *SELL* signaling network (bottom). **(E)** Circle plot shows the inferred intercellular communication network for *Thbs* signaling highlighting the autocrine and paracrine signaling to immune and endothelial cells (top). Heatmap shows the relative importance of each cell group based on the computed four network centrality measures of *Thbs* signaling network (bottom). **(F)** Heatmap showing the differential number of interactions and interaction strength in the male and female PND 1 lung. The top colored bar plot represents incoming signaling, while the right colored bar plot represents outgoing signaling. Pathways highlighted in blue (or purple) represent increased (or decreased) signaling in males compared to females. **(G)** Stacked bar graph showing the significant signaling pathways ranked based on differences in the overall information flow within the inferred networks between male and female lung. The top signaling pathways colored purple are enriched in the female lung and those colored blue were enriched in the male lung. **(H)** Circle plot shows the inferred intercellular communication network for *Tnf* (in female lung) and **(I)** *Tweak* (in male lung) signaling highlighting the autocrine and paracrine signaling to immune and endothelial cells. Solid and open circles represent source and target, respectively. Circle sizes are proportional to the number of cells in each cell group and edge width represents the communication probability. Edge colors are consistent with the signaling source. Heatmap shows the relative importance of each cell group based on the computed four network centrality measures of *Tnf and Tweak* signaling network.

Network centrality analysis of the *Angptl* signaling identified that alveolar and transient macrophages are the most prominent sources and dominant mediators for this ligand and has a predominant paracrine effect on other endothelial cells (**Figure 10B**). As described previously, *Angptl4* is a potent angiogenic factor [61] and has anti-inflammatory activity [62,63]. The effects of *Angptl4* could be mediated through integrin or syndecan receptors on the endothelial cells. Conversely, a signaling molecule that was sourced from endothelial cells and targeting alveolar and transient macrophages was CD200 (**Figure 10C**), which is a highly glycosylated cell surface protein. The CD200/CD200R immune ligand-receptor system decreases tissue resident macrophage activation [132–137]. The SELL (L-Selectin) signaling pathway was identified between the classical monocytes, neutrophils (immune cells) as sources and all the lung endothelial cells as targets. L-selectin is a major regulator of leukocyte adhesion, migration and signaling (**Figure 10D**)[138]. Another interesting immune-endothelial interaction pathway that was identified was *Thbs (*thrombospondin-1), being sourced from monocytes and neutrophils and targeting mainly the lung capillary endothelium (**Figure 10E**). Monocytes as sources of *Thbs* and endothelial cells as targets have been reported in prior studies[139,140] but the role in early lung development has not been studied.

We show differential number of interactions and interaction strength in the male and female PND 1 lung in greater detail in **Figure 10F**. The top-colored bar plot represents incoming signaling, while the right colored bar plot represents outgoing signaling. Values highlighted in blue (or purple) represent increased (or decreased) signaling in males compared to females. Interestingly, signaling from NK cells (both incoming and outgoing) is predominant in females. The incoming (increased in males) and outgoing (increased in females) signaling in venous endothelial cells varied by sex. In the alveolar macrophages, both the incoming and outgoing signaling was increased in females. The interaction strength of outgoing and incoming signaling pathways among classical monocytes and neutrophils was increased in males.

Next, we wanted to identify the conserved and context-specific signaling pathways and compared the information flow in the male and female lung (**Figure 10G**). The top signaling pathways colored purple are enriched in the female lung, and blue pathways were enriched in the male lung. One of the female-specific highlighted pathways is tumor necrosis factor (**Figure 10H**). Alveolar macrophages are the predominant source with majority of other immune and endothelial cells being targets. The TWEAK signaling pathway was male-specific (**Figure 10I**), with alveolar macrophages being the sole target with lung endothelial cells (except aCaps) and other macrophages being sources in this signaling pathway. TWEAK is a TNF superfamily cytokine that regulates inflammatory responses and macrophages express the specific receptor for this cytokine [141,142].

### Novel endothelial-immune interactions in the alveolar stage of lung development postnatal lung day 21

**Figure 11** summarizes the cell-cell communication between the different endothelial and immune cell subpopulations at PND 21. We identified the global communication patterns that connect cell groups with signaling pathways in the context of outgoing or incoming signaling. The outgoing signaling shows seven patterns (**Supplemental Figure 8A**). The endothelial cells separated into gCaPs/arterial/venous (pattern 1), and aCaPs (pattern 5). Among the immune cells, neutrophils (pattern 7), alveolar macrophages (pattern 3), interstitial macrophages and dendritic cells (pattern 6), basophils (pattern 4) and B/NK/T/TReg cells (pattern 2) formed their respective outgoing patterns. The signaling molecules associated with these patterns are highlighted in **Supplemental Figure 8A**. The incoming signaling patterns grouped the endothelial cells; gCaP/arterial/venous (pattern #1), and the aCaPs (pattern #7) in three groups. Among the immune cells, interstitial macrophages, classical and non-classical monocytes (pattern #2), alveolar macrophages (pattern 5), neutrophils (pattern #8), basophils (pattern 6), dendritic cells (pattern 4), and T cells (pattern#3), comprised the six patterns. The signaling molecules potentially driving the patterns in target cells are included in **Supplemental Figure 8B**. The number and strength of interactions among and between endothelial and immune cells is shown in **Figure 11A**. The lung endothelial cells show a robust communication pattern with autocrine signaling among themselves. Interestingly, the number of interactions and the interaction strength of lung immune cells is greater with endothelial cells than among other immune cells, pointing to the role of these intercellular communications patterns during the alveolar stage of lung development.

**Figure 11:**
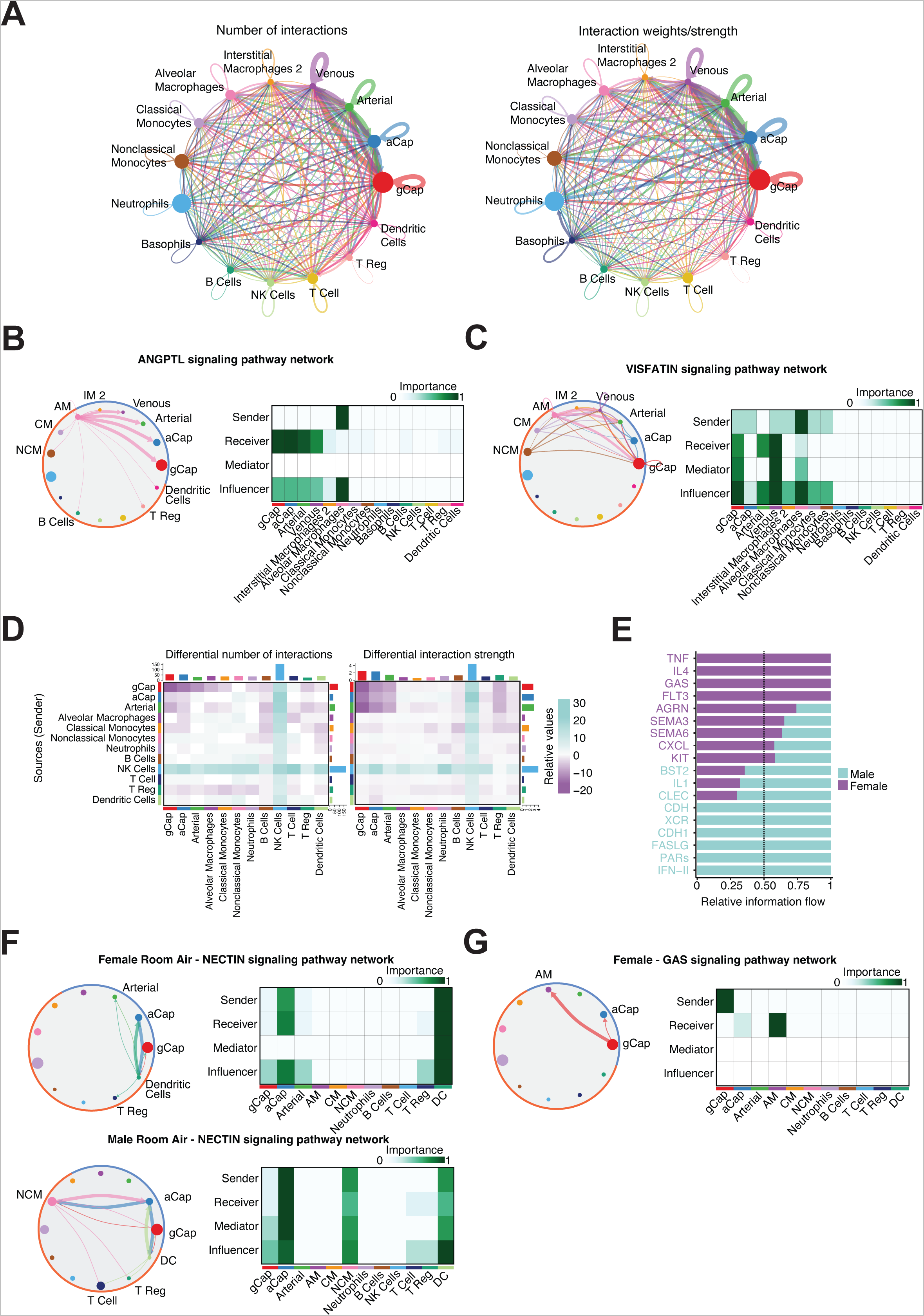
Novel endothelial-immune interactions in the postnatal lung at PND 21. **(A)** Circle plot showing the number of interactions between endothelial and immune cells. Edge colors are consistent with the sources as sender, and edge weights are proportional to the interaction strength. Thicker edge line indicates a stronger signal. Circle sizes are proportional to the number of cells in each cell group. **(B)** Circle plot and heatmap shows the inferred intercellular communication network for *Angptl* and **(C)** *Visfatin* signaling highlighting the autocrine and paracrine signaling to immune and endothelial cells. Circle sizes are proportional to the number of cells in each cell group and edge width represents the communication probability. Edge colors are consistent with the signaling source. Heatmap shows the relative importance of each cell group based on the computed four network centrality measures of *Angptl* and *Visfatin* signaling network. **(D)** Heatmap showing the differential number of interactions and interaction strength in the male and female PND 21 lung. The top colored bar plot represents incoming signaling, while the right colored bar plot represents outgoing signaling. Pathways highlighted in blue (or purple) represent increased (or decreased) signaling in males compared to females. **(E)** Stacked bar graph showing the significant signaling pathways ranked based on differences in the overall information flow within the inferred networks between male and female lung. The top signaling pathways colored purple are enriched in the female lung, and those colored blue were enriched in the male lung. **(F)** Circle plot and heatmap shows the inferred intercellular communication network for *Nectin* in male and female lung and **(G)** *Gas* signaling pathway network in the female lung, highlighting the autocrine and paracrine signaling to immune and endothelial cells. Circle sizes are proportional to the number of cells in each cell group and edge width represents the communication probability. Edge colors are consistent with the signaling source. Heatmap shows the relative importance of each cell group based on the computed four network centrality measures.

The *Angptl* signaling (**Figure 11B**) pathway was identified at PND 21 with the alveolar macrophages as the source and the lung endothelial cells as receivers. The *Visfatin* pathway is depicted in **Figure 11C**, with alveolar macrophages as the main source and venous, arterial and gCaPs being the receivers. *Visfatin*, also known as nicotinamide phosphoribosyltransferase (NAMPT), is an adipokine that was initially identified as a protein involved in nicotinamide adenine dinucleotide (NAD+) biosynthesis. *Visfatin* increased endothelial cell proliferation, migration, and tube formation by activating various signaling pathways involved in endothelial cell growth and survival [143–145] and increased nitric oxide production[146].

Next, we show differential number of interactions and interaction strength in the male and female PND 21 lung in greater detail in **Figure 11D**. The top colored bar plot represents incoming signaling, while the right colored bar plot represents outgoing signaling. Pathways highlighted in blue (or purple) represent increased (or decreased) signaling in males compared to females. Interestingly, at PND 21, signaling from NK cells (both incoming and outgoing) is predominant in males and dendritic cells in females. The signaling between the lung endothelial cells aCap, gCap, and arterial was increased in females. Shown in **Figure 11E** are signaling pathways enriched in female colored purple and enriched in male colored blue. We then selected a couple of pathways to highlight sex-specific differences. The Nectin signaling pathway is shown in **Figure 11F** where male non-classical monocytes secrete Nectin-1 but not females, with aCap cells being targets. Nectins belong to a family of cell adhesion molecules and play a role on vascular development and endothelial barrier function [147]. *Gas6* was a female-specific signaling pathway (**Figure 11G**). The *Gas6* gene encodes a protein called growth arrest-specific 6. gCaps serve as the source and alveolar macrophages as the main target. Gas6 can influence the activity of immune cells such as macrophages and can regulate the production of inflammatory cytokines and chemokines, and the balance between pro-inflammatory and anti-inflammatory responses [148–150]. Receptors for Gas6 *Axl* and *Mertk* are highly expressed in alveolar macrophages[151,152].

### Early hyperoxia exposure alters the cell-cell communication patterns between immune and endothelial cells at PND 21

Next, we show the differential number of interactions and interaction strength in the hyperoxia-exposed and control lung at PND 21 in **Figure 12A**. The top colored bar plot represents incoming signaling, while the right colored bar plot represents outgoing signaling. Pathways highlighted in red (or blue) represent increased (or decreased) signaling in hyperoxia compared to room air. Both incoming and outgoing signaling from the alveolar macrophages to the lung endothelial cells are increased in hyperoxia. In contrast, incoming and outgoing signaling in interstitial macrophages and NK cells are decreased in hyperoxia compared to room air. **Supplemental Figure 9** summarizes the cell-cell communication between the different endothelial and immune cell subpopulations in the hyperoxia-exposed lung at PND 21. The differential number of interactions and interaction strength in the male and female hyperoxia-exposed (PND 1-5) and control lung at PND 21 is shown in **Figure 12B**. Pathways highlighted in blue (or purple) represent increased (or decreased) signaling in male compared to female. Signaling to- and from-dendritic cells and NK cells was increased in females. Signaling between alveolar macrophages and lung endothelial cells was more pronounced in the male lung. One hyperoxia-specific effect was observed in the *Visfatin* (*Nampt*) pathway with loss of signaling from non-classical monocytes upon exposure to hyperoxia (**Figure 12C**). *Hspg2* or Perlecan is an endothelial cell-derived regulator of vascular homeostasis and is also a marker of endothelial injury [153–157]. Interestingly, in our study, venous endothelial cells were the predominant source of this signaling molecule with the other endothelial cells being the targets (**Figure 12C, top**). Venous endothelial cells induced *Hspg2* expression only upon exposure to hyperoxia (**Figure 12C, bottom**). The receptor *Dag1* was expressed in all endothelial cell sub-types.

**Figure 12:**
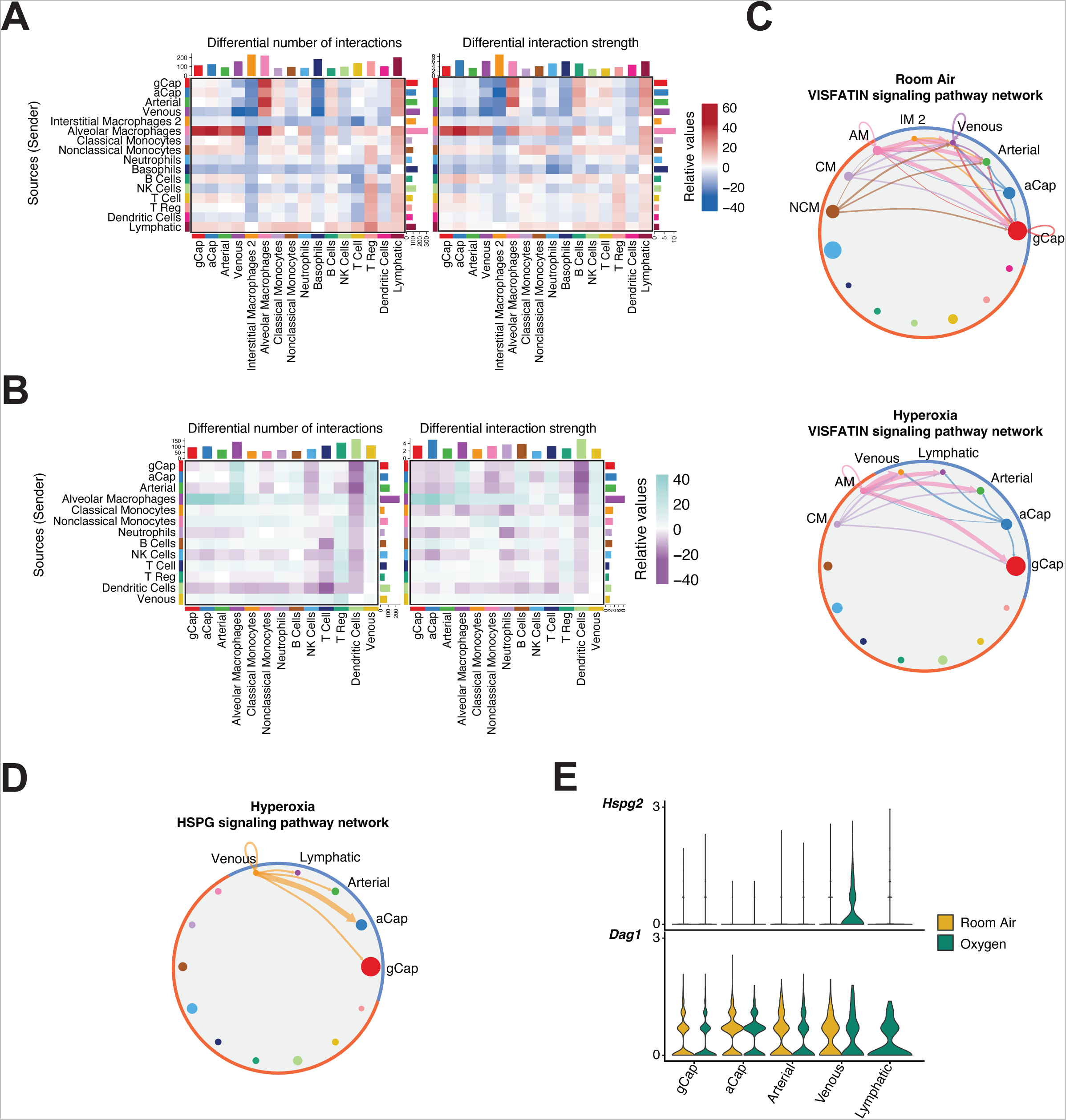
Early hyperoxia exposure alters the cell-cell communication patterns between the immune and endothelial cells. **(A)** Heatmap showing the differential number of interactions and interaction strength in the hyperoxia-exposed (95% FiO2;PND1-5) and room-air exposed murine lung at PND 21. The top colored bar plot represents incoming signaling, while the right colored bar plot represents outgoing signaling. Pathways highlighted in red (or blue) represent increased (or decreased) signaling in hyperoxia compared to room air. **B)** Heatmap showing the differential number of interactions and interaction strength in the hyperoxia-exposed (95% FiO2;PND1-5) male and female murine lung at PND 21. The top colored bar plot represents incoming signaling, while the right colored bar plot represents outgoing signaling. Pathways highlighted in blue (or purple) represent increased (or decreased) signaling in male compared to female. **(C)** Circle plot showing inferred intercellular communication network for *Visfatin* signaling network in room air (top)and hyperoxia (bottom). Different colors in the circle plot represent different cell groups. **(D)** Circle plot shows the inferred intercellular communication network for *Hspg2* signaling highlighting the autocrine and paracrine signaling to lung endothelial cells. Circle sizes are proportional to the number of cells in each cell group and edge width represents the communication probability. Edge colors are consistent with the signaling source. **(E)** Violin plots showing the expression of *Hspg2* in hyperoxia-exposed venous endothelial cells and the *Dag1* receptor expression in other endothelial cells.

## DISCUSSION

Investigating sex differences and their underlying mechanisms in lung diseases holds significant implications for human health. Gaining insights into why one sex may be more susceptible or resilient to disease pathogenesis identifies novel avenues for targeted treatments that can be beneficial for both sexes. Bronchopulmonary dysplasia (BPD) is a severe chronic lung disease that develops in preterm infants and has a strong bias for male sex. We previously used single-cell transcriptomics in the male and female neonatal murine lung acutely after hyperoxia exposure (95% FiO2; PND 1-PND 5; saccular stage of lung development) at PND 7 and reported the marked sex-specific changes in the lung immune, endothelial, and epithelial cell sub-populations. In this study, we highlight the sex-specific differences in individual lung cell sub populations at single-cell resolution immediately after birth at PND 1 and after early hyperoxic injury at PND 21 (alveolar stage of lung development). Our studies highlight the sex-specific differences immediately after birth in the PND 1 lung and describe the dynamic changes in the transcriptome of the endothelial and immune cells over the first three weeks of life, both in the unperturbed state and when the developing lung is exposed to early hyperoxic injury. We show that many hyperoxia-mediated effects on the murine lung are more pronounced immediately after injury and that differences in gene expression are much less pronounced at PND 21, however biological sex played an important role with more differentially expressed genes being sex-specific than common in both male and female lungs. Using pseudotime trajectory analysis, we highlighted gene clusters across the developmental windows and how these are altered by hyperoxia. Finally, we show that intercellular communication between the endothelial and immune cells at saccular and alveolar stages of lung development changes with sex-based biases in the crosstalk. Taken together, these data highlight the remarkable evolution of endothelial and immune cell transcriptional states across development and the differential sensitivity of the male and female lungs to hyperoxic injury.

Taking advantage of our previously published single-cell study of the PND 7 murine lung, we track the temporal differences in gene expression across cell types in the male and female lung. We show that alveolar macrophages are transcriptionally distinct in the male and female lung at PND 1, B cells are at PND 7, and T cells at PND 21. We also describe the *Mki67+ Top2a+* proliferating endothelial cells at PND 1 and PND 7 and show that these cells are transcriptionally more alike gCaps than aCaps with similar findings as reported by Zanini *et al* [34]. These cells are different from the proliferating endothelial cells that are present in the injured lung that express aCap markers as well [23,158]. Transient macrophages (*Galanin+*) were also identified in the early postnatal lung at PND 1 and not at the subsequent time-points. This unique cluster has been described in a recent publication[33] by Domingo-Gonzalez *et al.*, where before birth, they were identified around developing pulmonary vessels with more widespread distribution at PND 1. Trajectory analysis revealed that these macrophages give rise to alveolar macrophages during later lung development. The developmental role of the transient macrophages in early lung development and its role in the establishment of the alveolar macrophage niche needs to be investigated in future studies. Interestingly, transient macrophages from the female lung enriched for TNF-alpha signaling, which as an autocrine factor facilitates macrophage differentiation [159]. We also report the relative changes in the lung interstitial macrophage (IM) population across the saccular and alveolar phases of lung development and its alterations with early hyperoxia exposure. The IM1 and IM2 subsets of interstitial macrophages correspond to the CD206+ that localize in the per-bronchial regions and around vessels, secrete immunoregulatory cytokines and restrain tissue fibrosis and CD206-that localize in the alveolar interstitium and have antigen-presenting roles [37,96,160] respectively as described before. Interestingly, there is an increase in the IM2 and a decrease in the IM1 with lung development and hyperoxia decelerates this change. Lastly, at PND 1, we identified a “neutrophil-like” classical monocyte like cluster thought to arise from granulocyte-monocyte progenitors with immunoregulatory functions [64,65].

Sex-specific differences between male and female neonatal lung at PND 1 and PND 21 highlighted shared pathways that were distinct between the sexes. The interferon pathway was enriched in the male lung, while the TNF-alpha signaling pathway via NF-kappaB was enriched in the female lung. The known roles of component genes within these pathways such as *Nr4a1, Zfp36*, and *Psmb8* have been highlighted in the previous sections of the manuscript. We speculate that the male and female neonatal lung may diverge along these biological pathways during the postnatal lung development. We have previously shown sex-specific differences in the NF-kappaB pathway activation in the neonatal lung upon exposure to hyperoxia, with female lungs showing greater activation [17] and other groups have established the relevance of this pathway for postnatal lung angiogenesis [161]. Sex differences in immune response have been reviewed in detail before [162], but the studies in the context of normal neonatal lung development are lacking. In this study we also highlight cell-cell communication pathways between endothelial and immune cells at saccular and alveolar stage of lung development. We identified the *Angptl4* signaling pathway both at PND 1 and PND 21 signaling from the alveolar macrophages to the endothelial cells as well as sex-specific pathways; TNF-alpha and growth arrest-specific 6 (*Gas6*) signaling in females. Trajectory analysis on the lung capillary endothelial and macrophages from PND 1-PND 21 reveals the careful orchestration of gene clusters that are induced or repressed at the right time for proper developmental maturation. Distinct clusters of pseudotime-dependent genes (exclusive to room air or hyperoxia) and their gene expression trends in the lung endothelial cells and macrophages were identified. Transient macrophages at PND 1 give rise to alveolar macrophages and the *Inhba* + alveolar macrophages are more pronounced in the hyperoxia-exposed lung.

Following hyperoxia exposure from PND 1-5 during the saccular stage of lung development, the lung recovers in terms of cellular composition and transcriptional state at PND21. We show that the differences in gene expression between room air and hyperoxia in lung endothelial and immune cells are decreased at PND 21 compared to PND 7 (immediately after hyperoxia exposure). However, sex as a biological variable continues to play an important role in modulating the cellular transcriptional state with more sex-specific than shared genes among lung endothelial and immune cells. Alveolar macrophages and T cells are predominantly female biased and B cells and monocytes are male-biased in terms of the number of differentially expressed genes in the hyperoxia-exposed lung compared to room air controls. Interestingly, following hyperoxia exposure, the interferon pathway is negatively enriched in the male lung. Also, most sex-specific genes were not present on the sex chromosomes. Gene targets with potential role in modulation of hyperoxic lung injury and linked to sex chromosomes include *Tsc22d3*, *Xist* and *Kdm6a.* Intercellular communication patterns highlighted the interactions between alveolar macrophages and endothelial cells in the hyperoxia exposed-lung and sex-specific differences (signaling from- and to-dendritic cells and NK cells increased in females).

We recognize the limitations of the present study. We used pooled lung cells from three biological replicates, and thus an interaction analysis between sex and hyperoxia exposure is not possible. Our study was biased towards the lung endothelial and immune cells and thus lacking the insight from other equally important lung cell subpopulations. A detailed study of changes in relative cellular composition of different cellular subsets in the developing lung both at baseline and after exposure to hyperoxia was not performed. The strengths of this study include the detailed report of sex-specific differences in the lung endothelial and immune cell subpopulations at single-cell resolution, trajectory analysis, and the change in cell-cell communication patterns in the neonatal lung upon exposure to hyperoxia. Our study adds to the previous knowledge and identifies novel interaction pathways between specific immune and endothelial cells that are ripe targets for further exploration.

In summary, these data highlight the cellular heterogeneity and changes in the cellular transcriptional state through the saccular and alveolar stages of lung development and the crucial role of sex as a biological variable. We underline the rich intercellular communications between endothelial and immune cells and identify new receptor-ligand interactions between specific cells, which may be crucial for normal lung development and homeostasis. We also show that after early hyperoxia exposure, in the absence of ongoing injury, the neonatal murine lung recovers but some biological pathways and cell sub-populations remain perturbed and show sex-specific differences. Our data provide a detailed framework that enables a more complete understanding of disruptions in endothelial and immune cell transcriptional state, developmental trajectories, and cell-cell interactions due to early injury and modulation by sex as a biological variable.

## Acknowledgements

This work was supported in part by grants from National Institutes of Health [R01-HL144775 and R01-HL146395 to KL]. We acknowledge Dr. Lukas Simon for his help with initial approaches for data analysis. We also acknowledge Dominique Armstrong from helping with the single-cell isolation. Some of the figures in the publication were created with biorender.com using a paid subscription license.

## Supplemental Figures

**Supplemental Figure 1:**
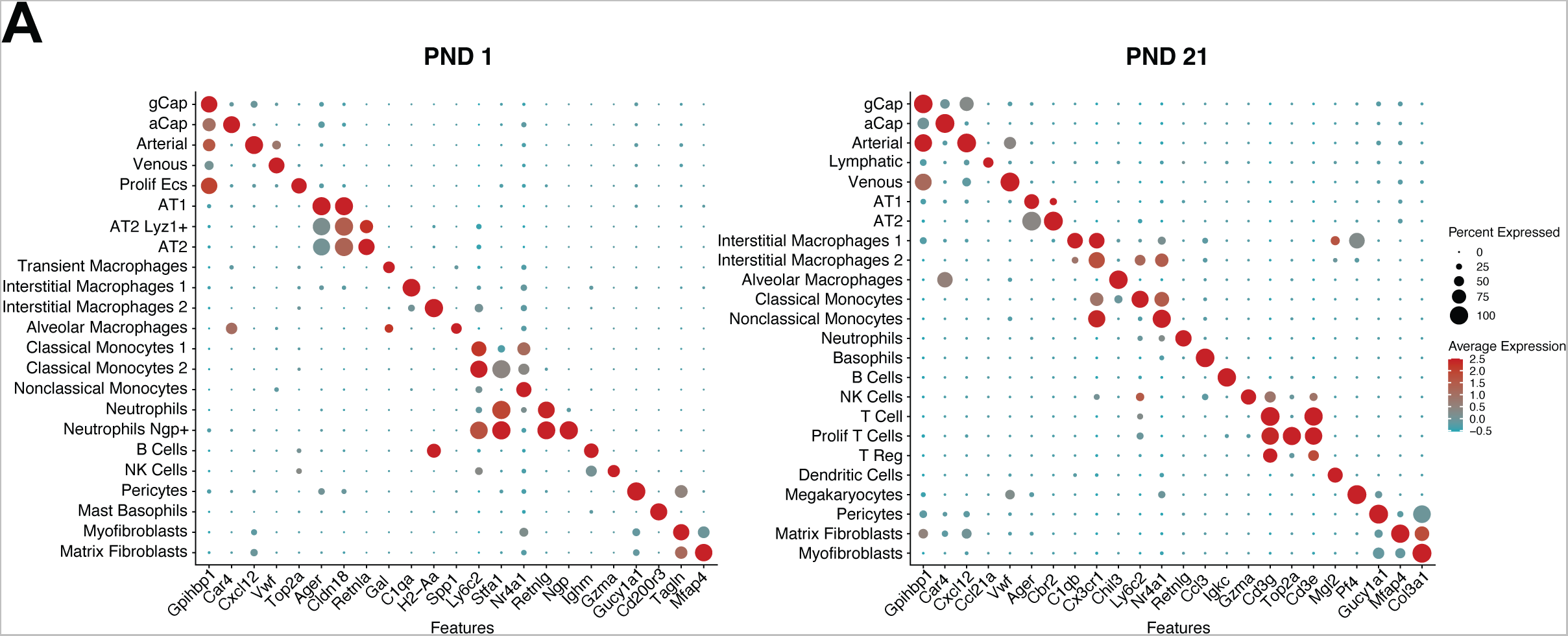
Dot plot showing expression levels (dot color) and percent of cells expressing each gene (dot size) of previously validated marker genes for each cluster at PND1 and PND21.

**Supplemental Figure 2:**
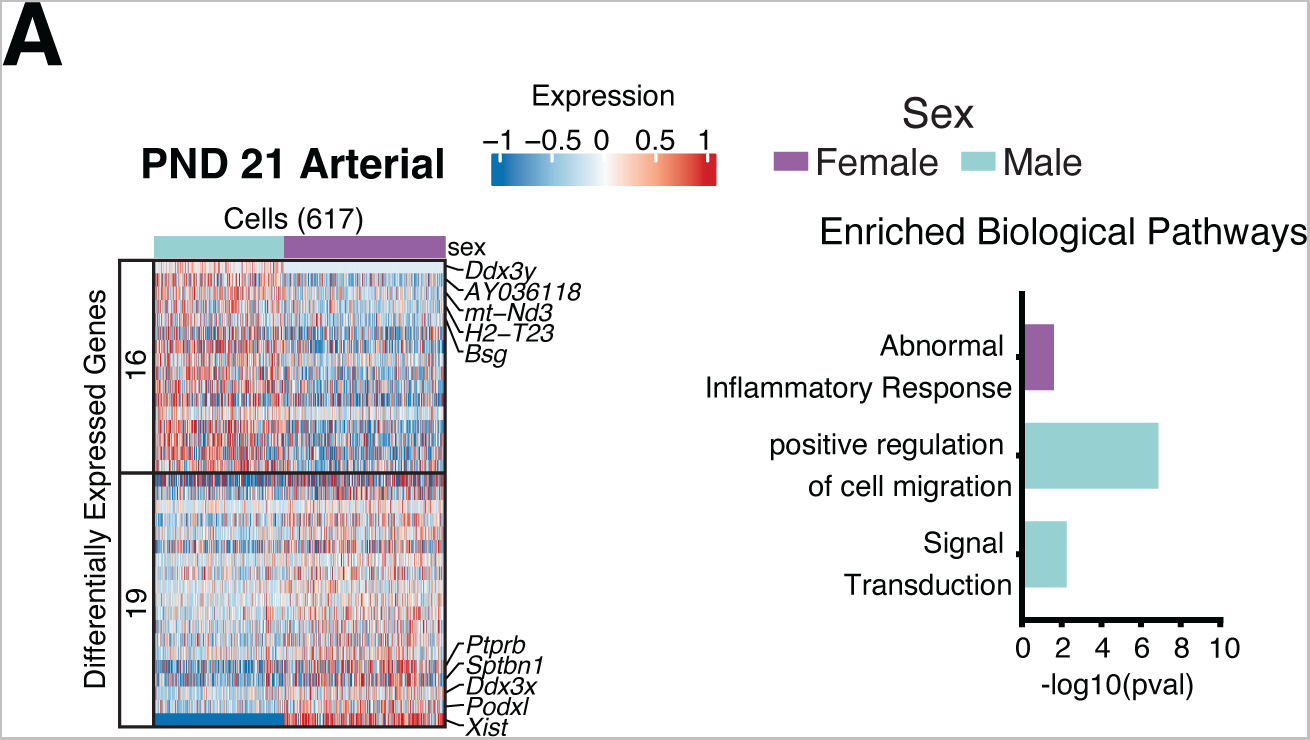
Sex-specific programming of biological pathways in the lung arterial endothelial cells during lung development. **(A)** Heatmap of differential expressed genes between male and female arterial endothelial cells at PND21 and enriched biological pathways. Number of cells sequenced are shown within parentheses.

**Supplemental Figure 3:**
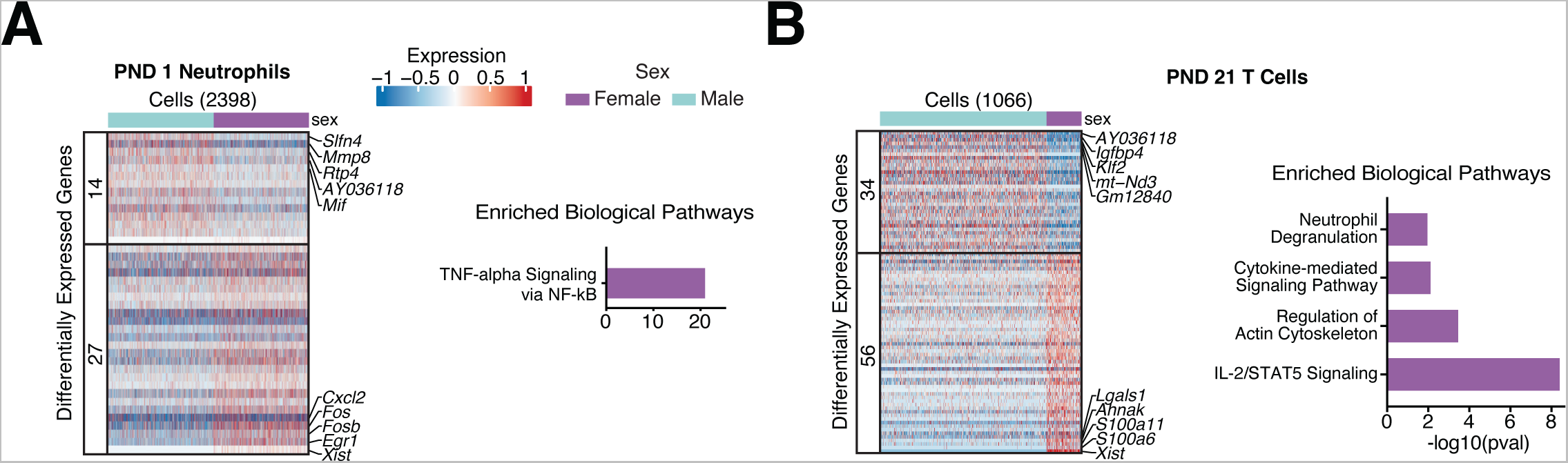
Sex-specific programming of biological pathways in the lung neutrophils and T cells during lung development. **(A)**. Heatmap of differentially expressed genes between male and female cells with enriched biological pathways in PND 1 neutrophils and **(B)** PND 21 T cells. Number of cells sequenced are shown within parentheses.

**Supplemental Figure 4:**
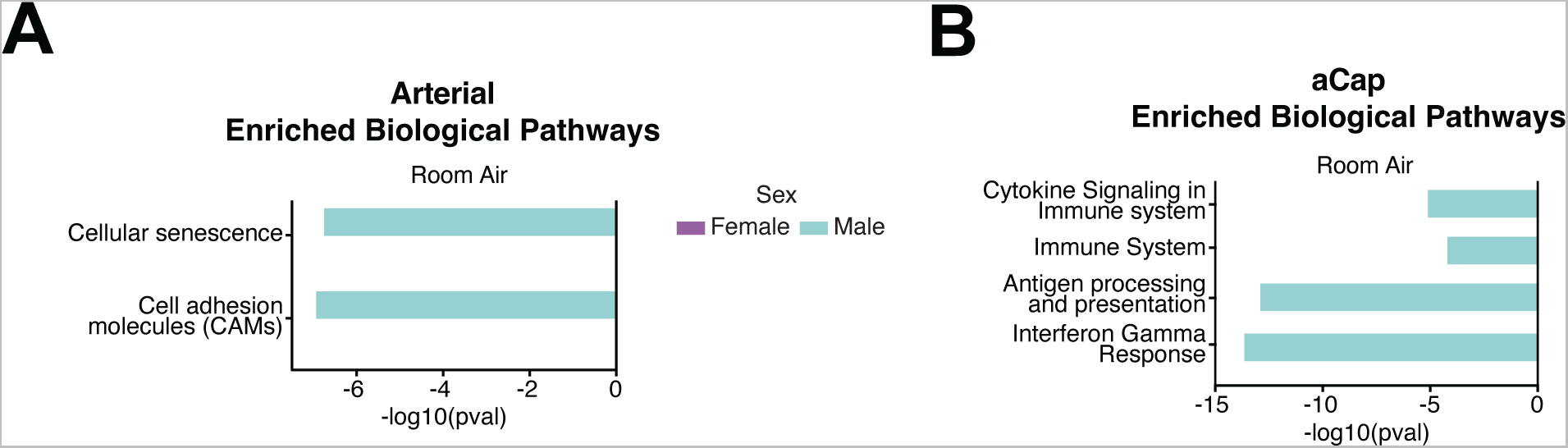
Enriched biological pathways in DEGs of arterial and aCap cells. **(A)**. Enriched biological pathways from DEGs in room air or hyperoxia at PND 21 arterial or **(B)** aCap cells.

**Supplemental Figure 5:**
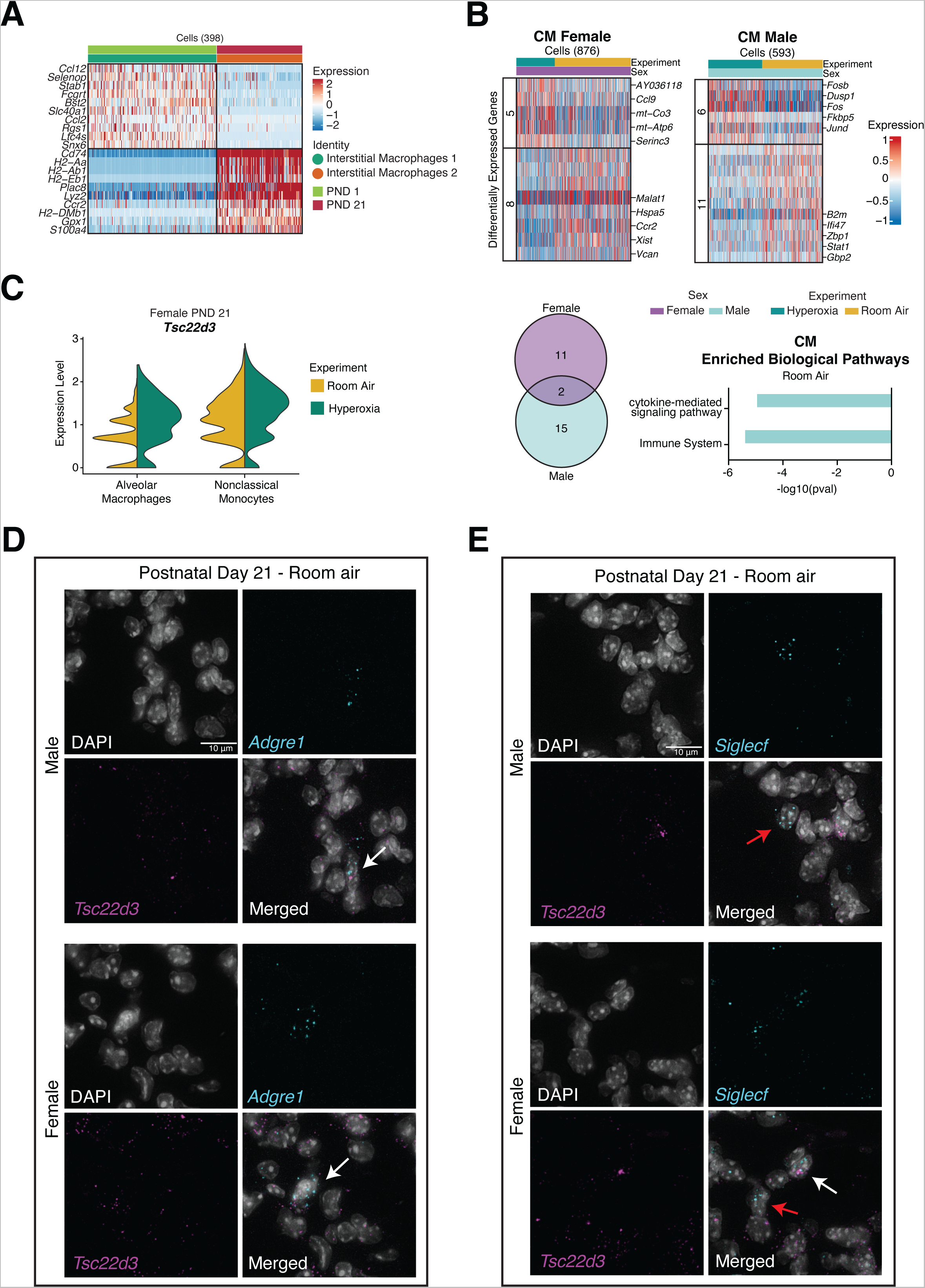
**(A)** Heatmap showing markers unique to interstitial macrophages 1 and interstitial macrophages 2. **(B)** Heatmap showing differentially expressed genes between room air and hyperoxia in female and male, Venn diagram showing the number of unique (sex-specific) and common DEGs in hyperoxia-exposed lungs and enriched biological pathways in classical monocytes at PND21. **(C)** Violin plots showing *Tsc22d3* expression in female lung alveolar macrophages and non-classical monocytes in room air and hyperoxia at PND21 **(D)** *In situ* hybridization of *Tsc22d3* and *Adgre1* in room air PND 21 male and female lungs. White arrows pointing to *Tsc22d3*+ and *Adgre1*+ cells. **(E)** *In situ* hybridization of *Tsc22d3* and *Siglecf* in room air PND 21 male and female lungs. White arrows point to *Tsc22d3*+ and *Siglecf* + cells. Red arrows point to *Tsc22d3*- and *Siglecf* + cells.

**Supplemental Figure 6:**
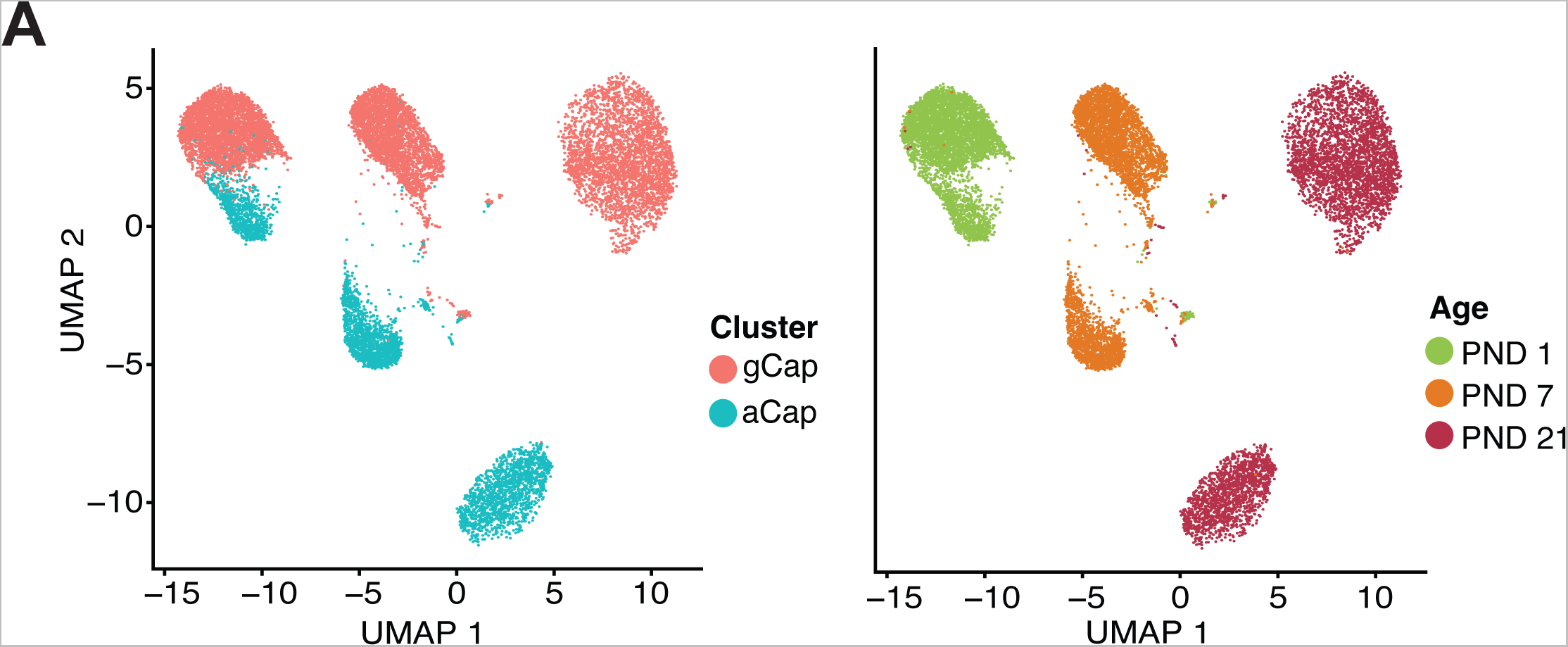
UMAP of sequenced lung aCaps and gCaps at PND 1, PND 7 and PND 21 split by cluster (left) and age (right).

**Supplemental Figure 7:**
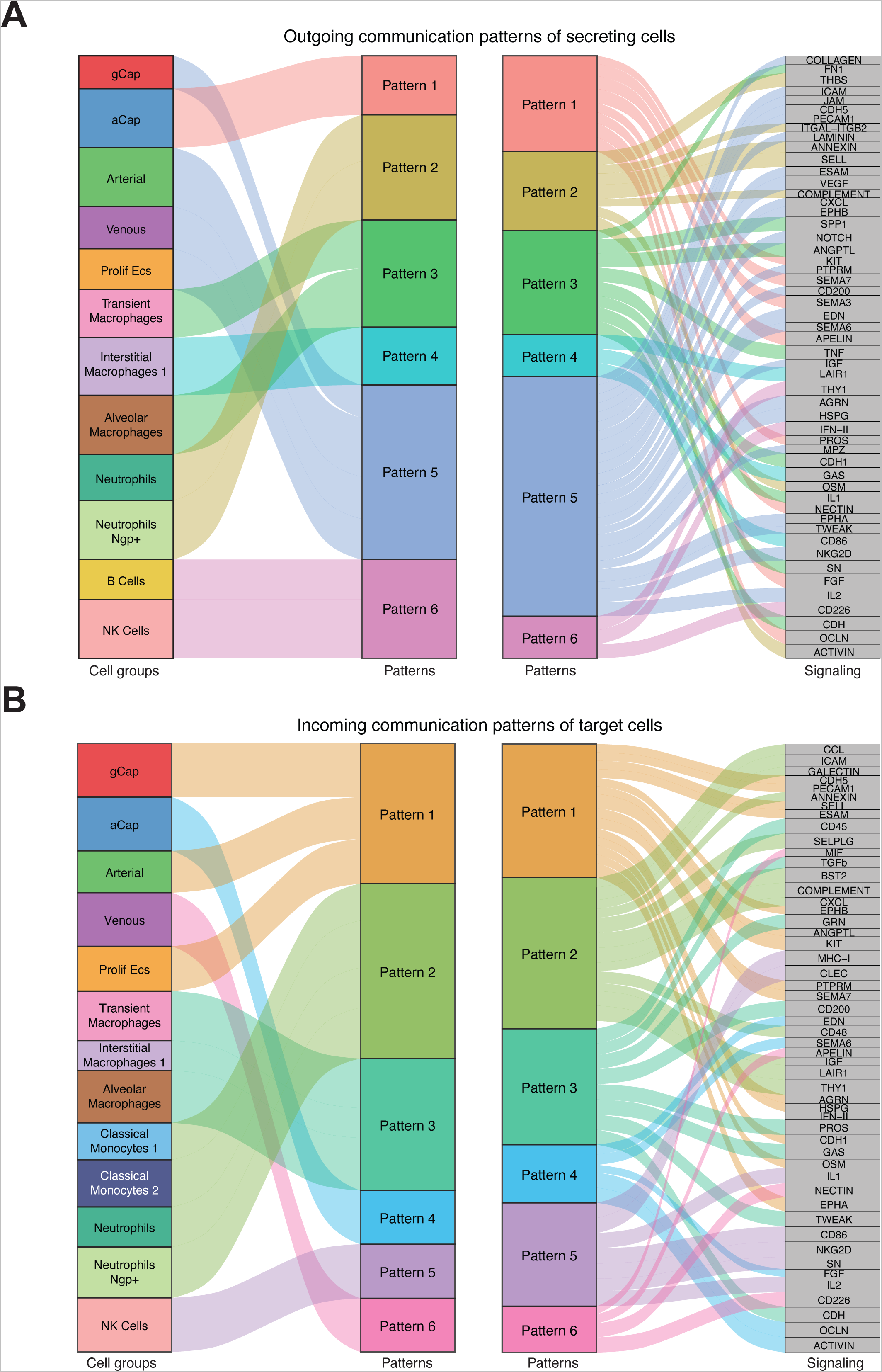
**(A)** Inferred outgoing and **(B)** incoming communication patterns of secreting cells and targets. The thickness of the flow indicates the contribution of the cell group or signaling pathway to each latent pattern.

**Supplemental Figure 8:**
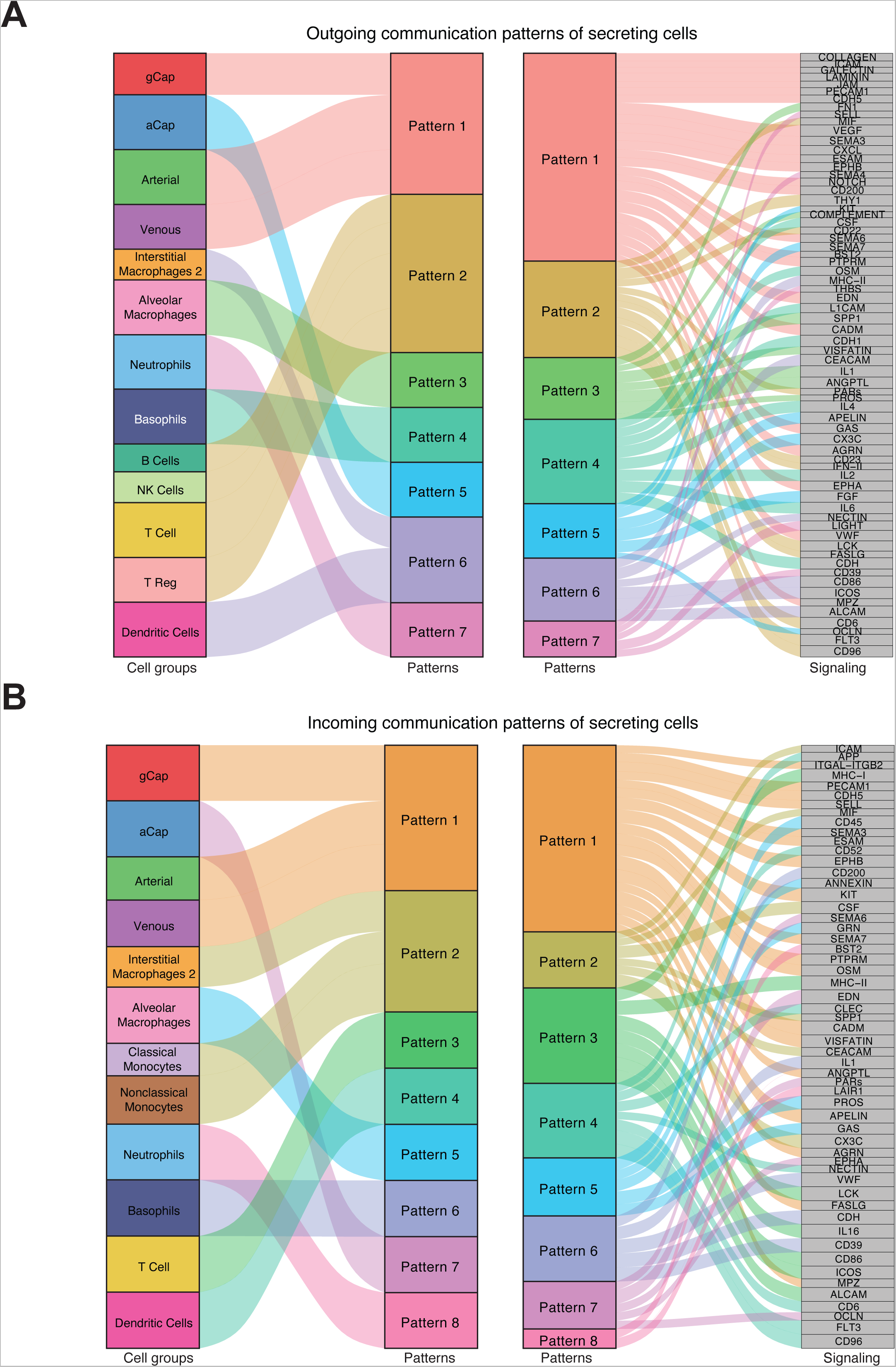
**(A)** Inferred outgoing and **(B)** incoming communication patterns of secreting cells and targets. The thickness of the flow indicates the contribution of the cell group or signaling pathway to each latent pattern.

**Supplemental Figure 9:**
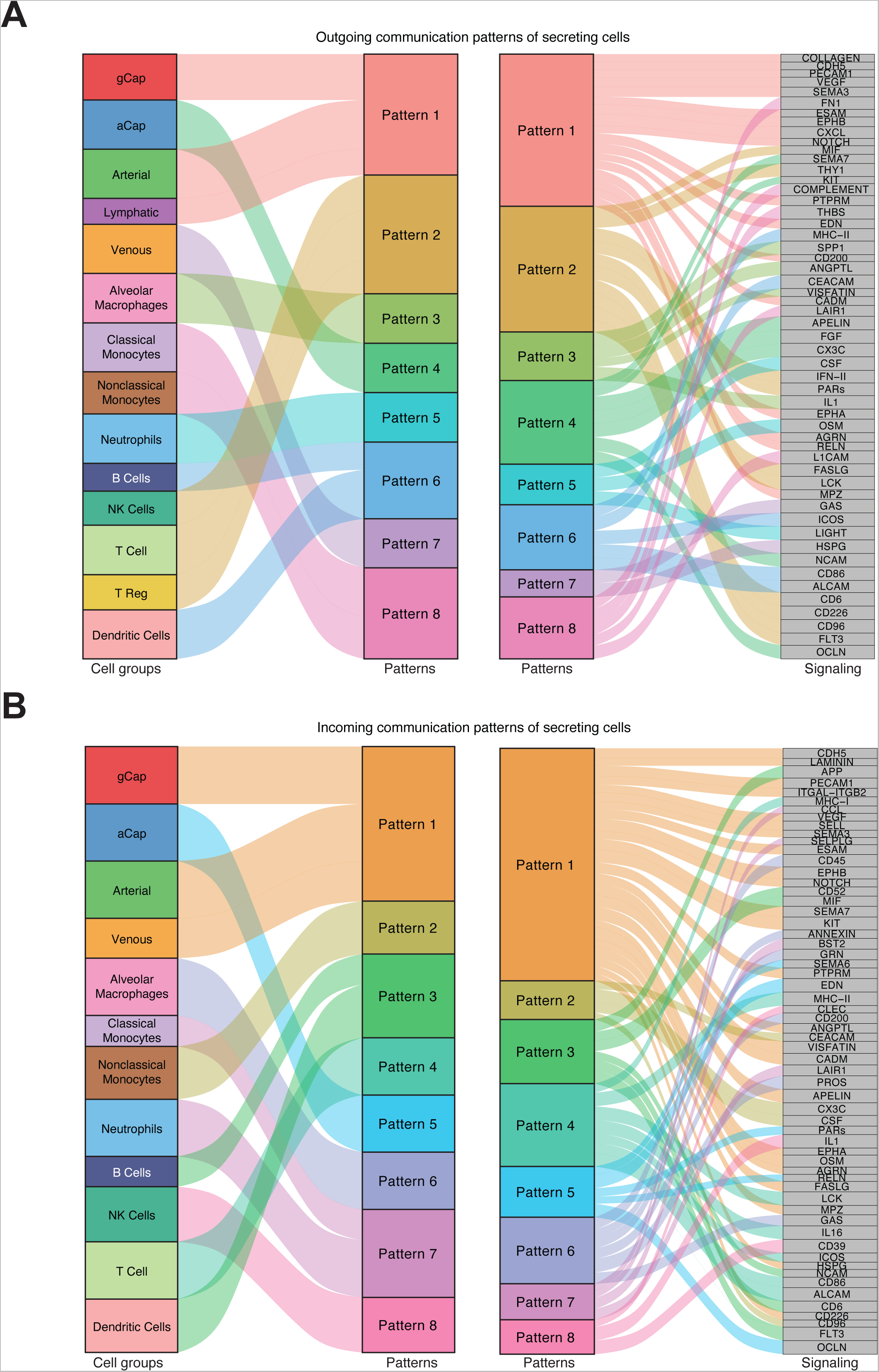
**(A)** Inferred outgoing and incoming communication patterns of secreting cells and targets. The thickness of the flow indicates the contribution of the cell group or signaling pathway to each latent pattern.

**Supplemental Table 1: Cluster gene markers for PND 1 lung cells.**

**Supplemental Table 2: Cluster gene markers for PND 21 lung cells.**

**Supplemental Table 3:** Differentially expressed genes in PND 1 and PND21 aCaps by sex and gene ontology enrichment analysis.

**Supplemental Table 4:** Differentially expressed genes in PND 1 and PND 21 gCaps by sex and gene ontology enrichment analysis.

**Supplemental Table 5:** Differentially expressed genes in PND 1 transient macrophages by sex and gene ontology enrichment analysis.

**Supplemental Table 6:** Differentially expressed genes in PND 1 classical monocytes 1 by sex and gene ontology enrichment analysis.

**Supplemental Table 7:** Differentially expressed genes in PND 1 alveolar macrophages by sex and gene ontology enrichment analysis.

**Supplemental Table 8:** Differentially expressed genes in PND 21 non classical monocytes by sex and gene ontology enrichment analysis.

**Supplemental Table 9:** Differentially expressed genes in PND 1 neutrophils by sex and gene ontology enrichment analysis.

**Supplemental Table 10:** Differentially expressed genes in PND 21 T cells by sex and gene ontology enrichment analysis.

**Supplemental Table 11:** Differentially expressed genes between room air and hyperoxia-exposed mice at PND 21 in endothelial and immune cells.

**Supplemental Table 12:** Differentially expressed genes, overlap or lack thereof between male and female lung gCap cells and enriched biological pathways in the male and female hyperoxia-exposed cells compared to room air controls.

**Supplemental Table 13:** Differentially expressed genes, overlap or lack thereof between male and female lung aCap cells and enriched biological pathways in the male and female hyperoxia-exposed cells compared to room air controls.

**Supplemental Table 14:** Differentially expressed genes, overlap or lack thereof between male and female lung arterial cells and enriched biological pathways in the male and female hyperoxia-exposed cells compared to room air controls.

**Supplemental Table 15:** Differential expression of genes between interstitial macrophages 1 and interstitial macrophages 2.

**Supplemental Table 16:** Differentially expressed genes, overlap or lack thereof between male and female lung alveolar macrophages and enriched biological pathways in the male and female hyperoxia-exposed cells compared to room air controls.

**Supplemental Table 17:** Differentially expressed genes, overlap or lack thereof between male and female lung non classical monocytes and enriched biological pathways in the male and female hyperoxia-exposed cells compared to room air controls.

**Supplemental Table 18:** Differentially expressed genes, overlap or lack thereof between male and female lung neutrophils and enriched biological pathways in the male and female hyperoxia-exposed cells compared to room air controls.

**Supplemental Table 19:** Differentially expressed genes, overlap or lack thereof between male and female lung classical monocytes and enriched biological pathways in the male and female hyperoxia-exposed cells compared to room air controls.

**Supplemental Table 20:** Other software used

